# Detection of pathways affected by positive selection in primate lineages ancestral to humans

**DOI:** 10.1101/044941

**Authors:** J.T. Daub, S. Moretti, I. I. Davidov, L. Excoffier, M. Robinson-Rechavi

**Affiliations:** CMPG, Institute of Ecology and Evolution, University of Berne, Baltzerstrasse 6, 3012 Berne, Switzerland; SIB Swiss Institute of Bioinformatics, 1015 Lausanne Switzerland; Dept. of Ecology and Evolution, University of Lausanne, 1015 Lausanne, Switzerland

## Abstract

Gene set enrichment approaches have been increasingly successful in finding signals of recent polygenic selection in the human genome. In this study, we aim at detecting biological pathways affected by positive selection in more ancient human evolutionary history. Focusing on four branches of the primate tree that lead to modern humans, we tested all available protein coding gene trees of the Primates clade for signals of adaptation in these branches, using the likelihood-based branch site test of positive selection. The results of these locus-specific tests were then used as input for a gene set enrichment test, where whole pathways are globally scored for a signal of positive selection, instead of focusing only on outlier “significant” genes. We identified signals of positive selection in several pathways that are mainly involved in immune response, sensory perception, metabolism, and energy production. These pathway-level results are highly significant, even though there is no functional enrichment when only focusing on top scoring genes. Interestingly, several gene sets are found significant at multiple levels in the phylogeny, but different genes are responsible for the selection signal in the different branches. This suggests that the same function has been optimized in different ways at different times in primate evolution.

## Introduction

An important challenge in the study of protein evolution is the detection of substitutions fixed by positive selection on a background of genetic drift and purifying selection. The detection of such positive selection signal has progressed thanks to better codon models and statistical tests (Delport et al. 2009). Yet these tests suffer from low power (Anisimova and Yang 2007; Gharib and Robinson-Rechavi 2013), especially when they are applied to closely related species, where phylogenetic trees only have relatively short branches with few descending sequences. The situation is worse when positive selection is weaker, and thus harder to detect, e.g. in species with small population sizes. If cumulated, these effects make it notably difficult to reliably detect positive selection in recent primate evolution, such as on the phylogenetic branches directly leading to humans.

Despite these inherent limitations, there is much interest in detecting positive selection in humans, their primate relatives, and their direct ancestors (Lachance and Tishkoff 2013). For example, as soon as the chimpanzee genome was available, genome-wide scans using codon models were performed, resulting in the detection of fast evolving genes related to immunity, host defense, or reproduction (Chimpanzee Sequencing and Analysis Consortium 2005; Nielsen et al. 2005). However, these genome scans often lacked power to distinguish positive selection from relaxed purifying selection. With more species (i.e., more data) positive selection was eventually detected. However, very few genes remained significant after correcting for multiple tests (e.g. Bakewell et al. 2007; Gibbs et al. 2007), and thus only those genes with many positively selected mutations were identified. While these limited results are interesting, recent studies have tried to uncover a more comprehensive picture of positive selection in the human lineage and its ancestors. Using patterns of incomplete lineage sorting (ILS), Munch et al. (2016) identified selective sweeps in ancestors of the Great Apes. Cagan et al. (2016) combined several neutrality tests to infer natural selection in the great apes, and found that population size has been a major determinant of the effectiveness of selective forces.

Here we propose to combine potentially weak to moderate signals from several genes to gain statistical power, using biologically meaningful groupings of genes, such as known regulatory and metabolic pathways. Indeed, several genes with small effect mutations can altogether have a large impact on a biological pathway, even though these genes would have little chance to be identified by conventional genome scans. An increasing number of studies has thus shifted the focus from single gene approaches to the detection of polygenic selection (e.g. Serra et al. 2011; Daub et al. 2013; Fraser 2013; Berg and Coop 2014; Carneiro et al. 2014; Wellenreuther and Hansson 2016), taking advantage of existing databases of gene sets and pathways. For example, we have used a gene set enrichment test to detect gene sets with polygenic adaptive signals in human populations (Daub et al. 2013), showing that most significant pathways for positive selection were involved in defense against pathogens. This procedure has also been successfully applied to find signals of convergent adaptation in humans living at high altitudes (Foll et al. 2014) or in tropical forests (Amorim et al. 2015) and to detect positive selection in ant genomes using the results of the branch-site test (Roux et al. 2014). More recently, we have extended the classical McDonald-Kreitman test (McDonald and Kreitman 1991) to detect more ancient selection signals, i.e. outlier pathways affected by different modes of selection after the split of humans with chimpanzee (Daub et al. 2015).

In the present study, we investigate whether one can find traces of positive selection in older periods of human evolution. We use the branch-site likelihood-based test, contrasting codon models with and without positive selection (Zhang et al. 2005). This testing procedure can detect episodic positive selection, while avoiding false positives due to relaxation of selective constraints, and it has been widely used to find signals of ancient positive selection in various species, including primates and other mammals (Zhang et al. 2005; Bakewell et al. 2007; Kosiol et al. 2008; Studer et al. 2008). We propose here to combine the gene-specific likelihoods of branch-site tests over all members of a gene set, to infer which biological systems have been under positive selection in the primate ancestors of humans.

Using this approach, we detect signals of adaptation in pathways involved in immune response, sensory perception, metabolism, and energy production. Furthermore, we find that in many candidate pathways different genes are responsible for the selective signals in different periods of primate evolution.

## Results

We have performed a gene set enrichment analysis to detect positive selection at the pathway level in the inner branches of a phylogenetic tree leading to African great apes (Homininae), great apes (Hominidae), apes (Hominoidae), and Old World monkeys and apes (Catarrhini) (Figure 1). We first ran separate branch-site tests of positive selection on inner branches of 15,738 protein coding gene trees of the Primates clade. The output of the procedure is a loglikelihood ratio test (ΔlnL) statistic, comparing the likelihood of a model without positive selection to that of a model with one additional parameter for positive selection. A branch in a gene tree with a high ΔlnL statistic reveals that a subset of codons is likely to have been positively selected (i.e. dN/dS>1) in that gene over that period.

**Figure 1:**
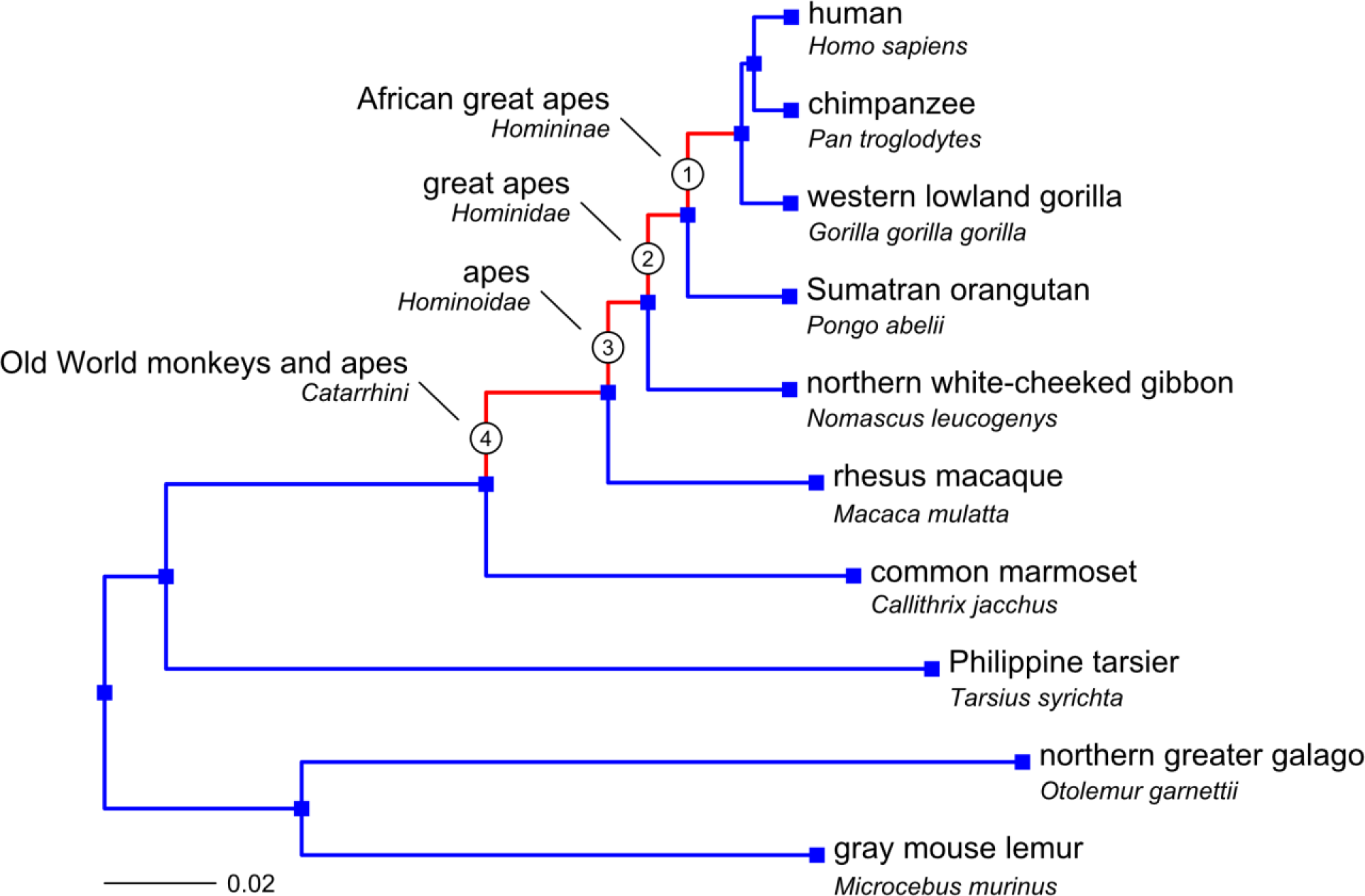
The Primates clade with the species used in the branch-site test. The four tested branches (Homoninae, Hominidae, Hominoidae, and Catarrhini) are numbered (used to identify branch specific lists of genes or gene sets, e.g. G1, G2, G3, G4) and marked in red. (Modified from the Ensembl mammalian species tree: https://github.com/Ensembl/ensemblcompara/blob/release/70/scripts/pipeline/species_tree_blength.nh. For more information about the construction of phylogenetic trees in Ensembl and the calculation of branch lengths, see http://dec2013.archive.ensembl.org/info/genome/compara/index.html)

We tested over 1,400 pathways from the Biosystems database (Geer et al. 2010) for episodic positive selection (Table 1). For each pathway and each branch, we calculated the sum of the ΔlnL4 values of the genes in this set (where ΔlnL4 is the fourth root of ΔlnL, see Materials and Methods), and we inferred the significance of this ‘SUMSTAT’ score (Tintle et al. 2009) against a null distribution of random gene sets of the same size. As shown in a previous study on the human specific branch (Daub et al. 2015), we find that genes in gene sets tend to be more conserved in internal branches of the Primates tree than genes that are not included in any gene set. This pattern is most pronounced in the two youngest branches leading to Homininae and Hominidae (Supplemental Figure S1). To account for the fact that some genes in sets are more conserved, we created null distributions that explicitly reflect the expected genomic background by randomly sampling from the group of genes that are part of at least one gene set.

**Table 1.** Number of gene sets and genes part of sets in the four tested branches. For each branch the number of significant sets (q < 0.2) in the SUMSTAT gene set enrichment test is reported, both before and after removing overlapping genes (‘pruning’), as well as the number of significant sets before pruning that remain significant after removal of their highest scoring gene.

| branch | leading to<sup>a</sup> | # sets | # genes in sets | # significant sets |  |  |
| --- | --- | --- | --- | --- | --- | --- |
|  |  |  |  | before pruning | without top scoring gene | after pruning |
| 1 Homininae | African Apes (Hu, Ch, Go) | 1,415 | 7,600 | 8 | 6 | 2 |
| 2 Hominidae | Great Apes (Hu, Ch, Go, Or) | 1,424 | 7,849 | 34 | 32 | 7 |
| 3 Hominoidea | Apes (Hu, Ch, Go, Gi) | 1,441 | 8,016 | 43 | 42 | 6 |
| 4 Catarrhini | Apes & Old World Monkeys (Hu, Ch, Go, Gi, Ma) | 1,441 | 8,058 | 95 | 93 | 9 |

<sup>a</sup>: Hu: human, Ch: chimpanzee, Go: gorilla, Or: orangutan, Gi: gibbon, Ma: macaque

As we are essentially interested in finding gene sets that show a global shift in selection scores and not those gene sets that include a single or a few extremely significant genes, we repeated the procedure after excluding in each set the highest scoring gene. Only pathways that scored a q-value below 0.2 (thus allowing a 20% FDR) both before and after exclusion of the top significant gene were considered candidates for polygenic selection.

We found 6, 32, 42, and 93 such significant pathways in the Homininae, Hominidae, Hominoidae and Catarrhini branches respectively (Table 1 and Supplemental Table S1). The fact that we find more candidate pathways when we go further back in evolutionary history is not necessarily caused by a change of selective pressures, but could be due to an increased power to detect selection in the longer ancient branches (see Discussion). In all four tested branches, we found clusters of candidate pathways that share a considerable proportion of their genes and often have similar biological functions, partly due to the nature of our data source, which is an aggregation of multiple pathway databases (Supplemental Figures S2 to S5). We removed this redundancy with a ‘pruning’ method, by iteratively removing the genes of the top scoring set from the remaining sets and rerunning the testing procedure on these remaining sets. The pruning procedure considerably reduced the number of significant sets and yielded 2, 7, 6, and 9 independent candidate sets in the Homininae, Hominidae, Hominoidae and Catarrhini branches respectively (Table 2). Even though all high scoring pathways could be worth further investigation, those gene sets that remain significant after pruning are the pathways with the best evidence for direct action of positive selection.

**Table 2.** Results of the SUMSTAT gene set enrichment test. For each branch, only the pathways that score significant (q < 0.2) both before and after pruning (removal of overlapping genes) are listed. The SUMSTAT scores and gene set sizes that changed after pruning are shown in parentheses. Pathways which score significant on more than one branch are highlighted by the symbol ‘§’.

| **Homininae** | *SUMSTAT*<br>(post-pruning) | *size*<br>(post-pruning) | *q*<br>post-pruning |
| --- | --- | --- | --- |
| GPCR downstream signaling § | 125.10 | 645 | 0.0032 |
| Immunoregulatory interactions between a Lymphoid and a non-Lymphoid cell § | 19.20 (18.61) | 54 (53) | 0.0047 |
| **Hominidae** | *score* | *size* | *q*<br>post-pruning |
| Olfactory Signaling Pathway | 69.34 | 230 | $<10^{-5}$ |
| Immunoregulatory interactions between a Lymphoid and a non-Lymphoid cell § | 22.34 | 57 | $<10^{-5}$ |
| Metabolism of xenobiotics by cytochrome P450 § | 16.81 | 57 | 0.1422 |
| Oxidative phosphorylation (WikiPathways) | 14.43 | 46 | 0.1422 |
| Intestinal immune network for IgA production | 12.68 (11.68) | 40 (36) | 0.1932 |
| Fatty acid metabolism § | 13.84 | 47 | 0.1932 |
| Synthesis of bile acids and bile salts via 7alpha-hydroxycholesterol | 8.08 (7.16) | 20 (17) | 0.1932 |
| **Hominoidae** | *score* | *size* | *q*<br>post-pruning |
| Staphylococcus aureus infection § | 23.88 | 44 | $<10^{-5}$ |
| GPCR downstream signaling § | 155.59 (150.17) | 687 (680) | $<10^{-5}$ |
| Electron Transport Chain | 30.13 | 84 | 0.0010 |
| Complement cascade | 16.36 (9.30) | 29 (16) | 0.0319 |
| Metabolism of xenobiotics by cytochrome P450 § | 20.11 | 59 | 0.0319 |
| Immunoregulatory interactions between a Lymphoid and a non-Lymphoid cell § | 22.34 (16.92) | 58 (52) | 0.1226 |
| **Catarrhini** | *score* | *size* | *q*<br>post-pruning |
| Hematopoietic cell lineage | 40.12 | 79 | $<10^{-5}$ |
| Non-alcoholic fatty liver disease (NAFLD) | 48.94 (47.02) | 129 (124) | $<10^{-5}$ |
| Cytokine-cytokine receptor interaction | 95.90 (73.29) | 236 (199) | $<10^{-5}$ |
| Staphylococcus aureus infection § | 24.11 (20.48) | 43 (37) | 0.0025 |
| Chemical carcinogenesis | 28.27 (26.87) | 63 (62) | 0.0066 |
| Defensins | 17.02 (16.47) | 40 (37) | 0.0896 |
| Pancreatic secretion | 29.93 (28.45) | 85 (83) | 0.1177 |
| Fatty acid metabolism § | 18.41 | 48 | 0.1814 |
| NF-kB activation through FADD/RIP-1 pathway mediated by caspase-8 and -10 | 7.98 (6.62) | 12 (10) | 0.1898 |

The strongest candidate for positive selection in the Homininae branch is the *GPCR downstream signaling* pathway, which is also a top scoring candidate in the Hominoidae branch. G proteincoupled receptors (GPCRs) are membrane proteins that regulate the cellular response to external signals such as neurotransmitters and hormones, and they play an important role in vision, taste and smell (Rosenbaum et al. 2009). We found that 28 out of the 43 high scoring genes in this pathway (having a ΔlnL4>1) are genes coding for olfactory receptors, suggesting that its role in taste and smell has been a major driver for selection.

The second candidate is the *Immunoregulatory interactions between a Lymphoid and a non-Lymphoid cell* pathway, which contains receptors and cell adhesion molecules that are important in immune response regulation of lymphocytes. This pathway is also significant after pruning in the Hominidae and Hominoidae branches, and it scores high before pruning in the Catarrhini branch. Interestingly, the genes in this pathway that contribute most to the polygenic selection signal differ among branches (Figure 2), which suggests that while the same pathway has been under selection over a long period, different genes underwent adaptive changes at different times. This is a pattern that is shared by many of the significant pathways (Supplemental Figure S6).

**Figure 2:**
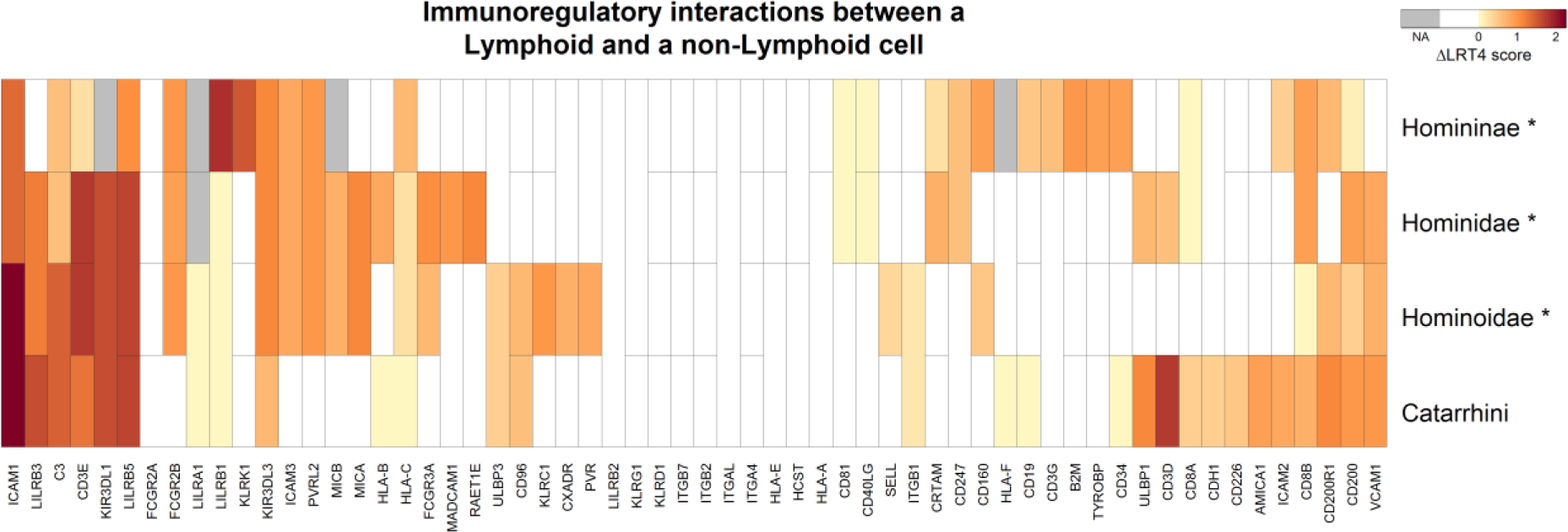
Heat map showing ΔlnL4 scores of genes in the *Immunoregulatory interactions between a Lymphoid and a non-Lymphoid cell* pathway for the four tested inner branches of the Primates tree. Branches where the pathway scores significant after pruning are marked with a ‘*’. The genes are grouped by hierarchical clustering to visualize blocks with similar signals within and among branches. Genes for which ΔlnL4 scores were not available (NA) in a certain branch are depicted in grey. Genes are merged (horizontally) with their paralog(s) into an ‘ancestral gene’ in the branches preceding a duplication and their scores were included only once in the calculation of the SUMSTAT score for these branches. Genes with (vertically) merged branches represent cases where the sequence of one or more species is missing or excluded, resulting in a single ‘average’ ΔlnL4 score over multiple branches. We use this score when testing each branch separately. The ΔlnL4 score is computed as the fourth root of log-likelihood ratio in the branch site test for positive selection.

The highest scoring candidate in the Hominidae branch is the *Olfactory Signaling Pathway*, i.e. smell perception. The pathway *Metabolism of xenobiotics by cytochrome P450*, which ranked third in Hominidae and fifth in Hominoidae, encodes detoxifying proteins which play a role in the metabolism of xenobiotics such as drugs and toxins; they include cytochrome P450 enzymes or glutathione S-transferases. Other candidate pathways are also involved in metabolism, such as the *Synthesis of bile acids and bile salts via 7alpha-hydroxycholesterol* and *Fatty acid metabolism*. Note that the latter pathway is also a candidate for positive selection in the Catarrhini branch. The remaining two top scoring pathways in the Hominidae branch are related to immune response (*Intestinal immune network for IgA production)* and electron transport (*Oxidative phosphorylation*).

In addition to the *Immunoregulatory interactions between a Lymphoid and a non-Lymphoid cell* pathway, two other immune response pathways remain significant after pruning in the Hominoidae branch. The highest scoring candidate, *Staphylococcus aureus infection*, includes genes coding for proteins used by the bacterium *S. aureus* to infect a host cell (Foster 2009). It is also a top candidate in the Catarrhini branch. The *Complement cascade* pathway plays an important role in immune surveillance and homeostasis (Ricklin et al. 2010). Finally, the last candidate in the Hominoidae branch, *Electron Transport Chain*, plays a role in energy production.

Five out of the nine pathways that remain significant after pruning in the Catarrhini branch have a function in immune response. Apart from the above-mentioned *Staphylococcus aureus infection*, these are *Cytokine-cytokine receptor interaction*, *Hematopoietic cell lineage* (giving rise to various types of blood cells including leukocytes), the host defense peptides *Defensins*, and *NF-kB activation through FADD/RIP-1 pathway mediated by caspase-8 and -10*. The fourth candidate is *Non-alcoholic fatty liver disease (NAFLD)*, a disease linked with metabolic syndrome (Dowman et al. 2010). Note that before pruning, this latter pathway was part of a cluster of gene sets related to electron transport, a process which – when disturbed – can lead to oxidative stress, which is a key feature of NAFLD pathology. The remaining candidate pathways have functions in the metabolism of toxins (*Chemical carcinogenesis*) and nutrients (*Pancreatic secretion*).

## Discussion

### Processes with evidence of positive selection

In this study, we have combined the strength of two approaches to detect positive selection in the ancestral lineages of humans. The branch-site test identifies episodes of selection at specific evolutionary times and sites in a protein, and gene set enrichment combines the signal of multiple genes to find selection at the pathway level. We identified several significant pathways related to immune response, sensory perception, metabolism, and electron transport in different branches of the primate tree (Supplemental Figures S2 to S5). These pathways were often organized in clusters that share many genes and have similar biological functions. After removal of the overlap between these pathways with a pruning procedure, two to nine pathways remained significant per branch. These pathways are our prime candidates for having been shaped by positive selection in primate evolution, and correspond to four broad biological processes.

First, in all four tested branches, immune response related pathways were among the top candidates, both before and after pruning. This strong and enduring signal suggests that the ongoing challenge of adaptation to changing pathogens has been one of the major selective pressures in primate evolution. This is in line with similar findings in previous reports of selection or fast evolution in ancient (Chimpanzee Sequencing and Analysis Consortium 2005; Nielsen et al. 2005) and recent human evolution (Daub et al. 2013).

Second, several significant pathways are involved in sensory perception, with *GPCR downstream signaling* and the *Olfactory Signaling Pathway* remaining after pruning in Homininae and Hominidae respectively. Sensory perception pathways have many functions, from sensing environmental signals to internal signals such as hormones and neurotransmitters. Earlier studies have reported genes involved in sensory perception to be evolving rapidly in humans and other primates, with some support for positive selection (Chimpanzee Sequencing and Analysis Consortium 2005; Nielsen et al. 2005; Arbiza et al. 2006; Kosiol et al. 2008). These genes could have been affected by positive selection because of changes in environment, behavior or diet.

Third, clusters of significant pathways involved in the metabolism of lipids and other nutrients were detected in Hominidae, Hominoidae and Catarrhini. This selective signal could be explained by changes in diet. Of note, the selective signal could also be due in part to the involvement of some of these pathways (*Metabolism of xenobiotics by cytochrome P450* and *Chemical carcinogenesis*) in the metabolism of potentially toxic xenobiotics. These results are specifically interesting, since adaptations in metabolism are expected with the changes in lifestyle that have marked hominid evolution, yet they have been rarely detected (but see Fumagalli et al. 2015; Mathieson et al. 2015). Our gene set approach thus allows us here to capture a subtle but biologically important signal of adaptation.

Fourth, there are several significant pathways involved in energy production in the Hominidae, Hominoidae and Catarrhini branches, with *Oxidative phosphorylation*, *Electron Transport Chain*, and *Non-alcoholic fatty liver disease (NAFLD)* remaining significant after pruning in these branches. Earlier candidate gene studies have also reported genes involved in electron transport to be under positive selection in anthropoid primates (reviewed in Grossman et al. 2004). It has been suggested that this could be related to anthropoid specific evolutionary changes, such as extended lifespan, prolonged fetal development and enlarged neocortex, as these traits require increased aerobic energy production (Grossman et al. 2004).

### Consistency of selection over evolutionary time

Several pathways reported here were also detected in an earlier study, in which we applied a gene set enrichment analysis on a more recent period of human evolutionary history (Daub et al. 2013). The *Cytokine-cytokine receptor interaction* pathway for example was detected as a potential candidate for recent polygenic human adaptation. Another candidate in that study, *Fatty Acid Beta oxidation,* scored significant here before pruning in Catarrhini, as did the *Malaria* pathway in all branches except Homininae. These findings suggest that certain functions have been under selection in multiple or ongoing episodes of positive selection, and might still be under selection now.

The finding of the *Olfactory Signaling Pathway* as candidate for positive selection is apparently in contradiction with a previous study (Daub et al. 2015) in which we found olfactory pathways to be under relaxed selection in the human branch. That earlier finding was in line with the fact that many olfactory genes have become pseudogenes in primates –and particularly in humans–, possibly because we have become more dependent on vision than smell and taste (Hughes et al. 2014). Our current results suggest that in more ancient Ape history, some of the olfactory system still played an important adaptive role, and evolved rapidly not just because of relaxed selection but because of adaptation. Two other pathways in the aforementioned study showed signs of positive selection, namely *Meiotic Recombination* and *Beta-catenin phosphorylation cascade*, but they did not score high in the current study. Again, this could be explained by positive selection acting differently over different time periods in Ape evolution. However, we cannot rule out that some differences in results are caused by the fact that we used different methods to test for selection in each study.

The fact that some pathways are significant over successive evolutionary periods (neighboring branches) appears to indicate that similar selective pressures occurred over a long timeframe in primates. Interestingly, we found that in such cases the highest scoring genes often differed among branches (Figure 2 and Supplemental Figure S6), which suggests that biological pathways under long-term selective pressures have adapted by means of changes in different genes over time, allowing the fine tuning of the pathway function without altering previous adaptations. However, we cannot exclude that we might lack power to detect more continuity in selective pressure in the same gene over long evolutionary periods.

We have also identified several candidate pathways that are branch-specific. Whereas these isolated adaptive signals may be due to unique changes in selective pressure, they might result from alternative causes. We therefore investigated if non-adaptive factors might have affected our results. We indeed find small but significant correlations between ΔlnL4 scores and gene tree size (number of branches; Pearson’s r = 0.07, p < 2.2e-16) or sequence length (r = -0.05, p < 2.2e-16). However, we found a more noticeable correlation between ΔlnL4 scores and gene specific branch length (estimated number of nucleotide substitutions per codon; r = 0.23, p < 2.2e-16), although the effect is reduced when we consider the synonymous branch length only (dS; r = 0.14, p < 2.2e-16). Finding a correlation between branch length and gene score makes sense since branch lengths are inferred from the number of mutations, and we have more power to detect positive selection with increasing numbers of mutations (Fletcher and Yang 2010), but positive selection can also lead to more mutations being fixed. So, our candidate pathways could be enriched in fast evolving genes, partly because positively selected genes evolve fast, and partly because there is more power to detect selection in fast evolving genes. Therefore, our finding that more significant pathways are observed in more ancient lineages might be partly due to an increased power to detect selection in the older branches that are on average longer than younger branches.

### Impact of GC bias and duplications

Earlier studies in primates have shown that GC-biased gene conversion can be confounded with positive selection as it leads to accelerated evolutionary rates and biased fixation of GC alleles in regions of high recombination (Berglund et al. 2009; Galtier et al. 2009; Ratnakumar et al. 2010). Although a simulation study by Gharib and Robinson-Rechavi (2013) indicated that the branch-site test is robust against GC variation, we have investigated a possible GC bias in our data by calculating correlations between GC content and selection score (ΔlnL4). We related selection scores with the GC content per branch, but also with the mean GC content per gene tree and with GC content variation (by calculating the GC content variance in a tree and the difference between the maximum and minimum GC content within a tree). We did not find any noticeable correlation between these statistics and selection scores (Supplemental Table S2A). Although one would not expect pathways to be GC-biased (but see Berna et al. 2012), we also measured correlations at the gene set level. We averaged the GC statistics per gene set and related them with the score (-log[p-value]) in the gene set enrichment test (Supplemental Table S2B). Again, we did not find any noticeable correlation with GC content, but we did detect a small correlation between GC variance (per gene tree) and gene set score (r between 0.15 and 0.19, depending on the tested branch, p-values all <1e-08). This can be attributed to the fact that fast evolving genes (gene trees with a relatively high number of mutations) will have on average a higher GC variance than slow evolving genes. That could explain why we see a higher correlation in gene sets than genes: in gene sets this effect is amplified, especially because genes in gene sets tend to have similar evolutionary regimes (Daub et al. 2015).

It has been hypothesized that duplicated genes can be a source of evolutionary novelties, through processes such as neo- or subfunctionalization or dosage effects (Innan and Kondrashov 2010). Indeed, Lorente-Galdos and colleagues showed recently that fast evolving exons in primates were enriched in duplicated regions (Lorente-Galdos et al. 2013). In another study, Qian and Zhang (2014) reported that simultaneously deleting a duplicate gene pair in budding yeast reduced fitness significantly more than deleting their singleton counterpart in fission yeast, again suggesting adaptation after duplication. We thus investigated the effect of duplications on our results. We find that branches with at least one duplication in their ancestral branches in the primate tree tend to have higher selection scores (i.e. 37% of these branches score ΔlnL>0, compared to 25% in branches without duplications in ancestral branches), suggesting an increase of selection intensity in some of the duplicated genes. Conversely, we applied the enrichment test on a reduced dataset containing only gene trees without duplications. With this new dataset we find less gene sets that are significant, which can be explained by a decrease in power as we have less genes and gene sets to test. Nevertheless, many of the significantly scoring gene sets from the original dataset still score significant or high with the new test, including the olfactory pathways that are known to contain many duplications (Supplemental Table S3). These new test results suggests that our results are not particularly confounded by duplications.

### Methodological limitations and strengths

A potential limitation of our study is that some genes are not included in the Selectome database, as only gene trees with at least six terminal leaves and passing stringent alignment quality filters were included (Moretti et al. 2014). Therefore, about 21% of genes were ignored in our enrichment test, which might lead to a deficit of trees with fast evolving genes that are difficult to align, as well as a potential excess of gene trees with duplications. The latter category contains several paralogs resulting in more leaves, and thus pass the criterion of six leaves even when a few sequences are missing from genomes or eliminated by alignment quality filters. To estimate the effect of this exclusion on our results, we tested for each gene set whether they had a significant excess of excluded genes (q-value < 0.2 with Fisher Exact test). We found 62 gene sets with such an excess (Supplemental Figure S7). They are mainly involved in protein metabolism (15 sets) and the cell cycle (37 sets). Interestingly, we also find a few gene sets that contain many excluded genes, but which still score high in our likelihood-based test, namely the *Olfactory signaling*, *Olfactory transduction* and *GPCR downstream signaling* pathways. Some of the pathways with an excess of excluded genes were candidates for being under selection in studies of recent human evolution, such as *Pathogenic Escherichia coli infection* (Daub et al. 2013), and we could thus not fully check if these pathways have also been under selection in more ancient primate evolution.

We want to emphasize that our gene set enrichment analysis differs from classical Gene Ontology (GO) enrichment tests, as we include all genes in our analysis instead of only considering top scoring genes after setting an arbitrary significance threshold. While GO enrichment tests ask whether these top scoring genes are enriched for a biological function or process (see e.g. Cagan et al. 2016), we aim to find pathways with an overall shift in the distribution of gene scores. For comparison, we also performed a GO enrichment test on the significant genes in each branch, but this procedure did not result in any significantly enriched GO terms. This negative result underlines the power of a gene set enrichment test as performed in this study, as we can detect pathways that contain many genes with small to moderate effect mutations.

### Conclusions

In conclusion, by combining the specificity of the branch-site test and the power of the gene set approach, we have been able to uncover for the first time strong signals of polygenic positive selection in several biological processes during long-term primate evolution. In addition to immune response, we find evidence for adaptive evolution on sensory perception, as well as on metabolism and energy production. The fact that different genes are involved in pathways showing signals of positive selection in several branches, suggests that the fine tuning of biological functions can change over time during primate evolution. Our results allow us to bridge the gap between studies of selection in deep mammalian evolution, and recent adaptation in the human lineage.

## Material and Methods

### Data collection

In this study we aim at detecting biological pathways affected by episodic positive selection in primate evolutionary history, specifically in the four inner branches of the Primates tree that lead to modern humans (Figure 1).

#### Branch-site likelihood test

The data used in this study was produced as part of release 6 of Selectome (http://selectome.unil.ch/, Proux et al. 2009; Moretti et al. 2014), which is a database that provides results of the branch-site likelihood test for positive selection (Zhang et al. 2005) on internal branches of several clades. The branch-site test can detect codon sites on specific phylogenetic branches that are affected by episodic positive selection. In short, it estimates the rate of non-synonymous (dN) and synonymous (dS) nucleotide substitutions to assess differences in selective pressure (dN/dS ratio) among branches and over sites. Usually, tested branches that have a class of sites with a dN/dS ratio ω_2_ > 1 are candidates for positive selection. While the strength of positive selection can in principle be estimated by the ratio ω_2_ or by the proportion p_2_ of sites in this class, we have found the likelihood ratio to be a good estimator of the evidence for positive selection (Studer et al. 2008; Roux et al. 2014). In more detail, for each branch, the maximum likelihood of the data is estimated under two models: one that allows for positive selection (H1), and one that only allows negative selection and neutral evolution (H0), and a loglikelihood ratio statistic ΔlnL = 2(lnLH1-lnLH0) is computed. To determine their significance, the ΔlnL values are usually compared to a chi-square (χ^2^) distribution with one degree of freedom; here we use all ΔlnL values without applying any *a priori* significance cut-off.

To avoid false positives due to poor sequence alignments, the Selectome pipeline includes many filtering and realignment steps to remove unreliable regions before running the branch site test (Moretti et al. 2014, and see http://selectome.unil.ch/cgi-bin/methods.cgi). Supplemental Figure S8 shows an example of a codon alignment before and after filtering, where filtering results in a significant change of the ΔlnL value. Furthermore, only the internal branches of gene families with at least six sequences (leaves in the tree) were computed in Selectome as there is more potential for errors on terminal branches, due to sensitivity to sequencing and annotation errors in one species; moreover, the test has low power and accuracy when only a few sequences are used (Anisimova et al. 2002).

For our gene set analysis, we thus used the ΔlnL values obtained by testing 15,738 gene trees from the Primates clade as defined in version 70 of Ensembl Compara (Vilella et al. 2009). We only kept test scores for the four branches mentioned above (and shown in Figure 1) if they led to at least one human gene. We could thus create for each branch *i* (Figure 1), a list *G_i_* with human genes and their corresponding ΔlnL scores. Because of duplication events and the lack of resolution of the Homininae label, these initial lists often contained several rows per gene, whereas we need at most one ΔlnL score per gene per branch for our enrichment test. On the other hand, missing or excluded sequences can result in branches lacking ΔlnL scores in a number of genes. We describe below how we dealt with these situations and how we handled computational issues with likelihood estimation.

#### Dealing with multiple Homininae branches

In Ensembl gene trees, both the branch that leads to the common ancestor of human, chimp and gorilla (Homininae) as well as the branch to the common ancestor of human and chimp (Hominini) are labeled as Homininae. As a result, many gene trees that have human, chimp and gorilla sequences present, have multiple branches annotated as Homininae. For example, the *DMXL1* gene family (Supplemental Figure S9) has a Homininae branch leading to the ancestor of human, chimp and gorilla followed by a branch leading to the ancestor of human and chimp which is labeled “Homininae”, although it is properly an Hominini branch. In about 42% of the gene trees we found such multiple Homininae labeled branches. In these cases we took the test scores of the oldest branch, which in 95% of the cases is also the longest branch.

#### Dealing with missing branches

For several genes the sequence of one or more species is absent or excluded due to low quality (Moretti et al. 2014). The corresponding branch in the gene tree is then merged with its downstream branch, and the ΔlnL score is assigned to this lower branch, while it actually represents an ‘average’ score over both branches. We therefore use this ΔlnL score for both branches as input in the gene set enrichment test. Supplemental Figure S10 shows an example of the C3 gene tree with a missing macaque sequence. Supplemental Table S4 shows results of the gene set enrichment test where in the case of a missing sequence, we assign the ΔlnL score only to the lower branch.

#### Dealing with gene duplications

Gene duplications will result in some species having paralogous genes. For our gene set enrichment analysis we removed the branches in the Primates tree that led to a duplication event (about 3% of all branches). Our procedure is further detailed in Supplemental Figure S11. Briefly, for branches predating the duplication, the values in the gene list corresponding to the duplicated genes, which are redundant (same ancestral branches reported for each paralog), were merged and replaced by one value, while for the branches after the duplication both values were kept, as each represents a different paralog.

#### Non-converging likelihoods

The numerical optimization (BFGS algorithm, see PAML documentation) does not always converge to the maximum likelihood estimates of H0 or H1. This can result in negative ΔlnL values or in false positives having extreme high ΔlnLs. In order to reduce the number of events of non-convergence, we ran the branch-site test two more times on the whole Primates tree, thus yielding a total of three likelihood scores for each of H0 and H1. We then selected for each branch the highest log-likelihoods for H0 and H1 among the three runs (lnLH0*_g_* and lnLH1*_g_*) and constructed the log-likelihood ratio score for a gene (ΔlnL*_g_*) as follows:

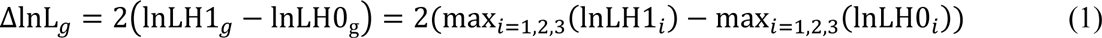

However, three runs could still allow for some non-convergence. If we obtained a negative ΔlnL*_g_* score (indication of non-convergence of all lnLH1 scores) we set ΔlnL*_g_* to zero (about 8% of the cases, with less than 0.06% having a ΔlnL*_g_* < -0.1).

The first branch-site test was run with *codeml* version 4.6 from PAML (Yang 2007). Most jobs of the first run were performed on the Swiss multi-scientific computing grid (SMSCG, http://lsds.hesge.ch/smscg/), while the longer jobs were submitted to the Vital-IT computer cluster (http://www.vital-it.ch) and to the Ubelix computer cluster of the university of Berne (http://www.ubelix.unibe.ch/). The second run was run with SlimCodeml (Schabauer et al. 2012), and the third with FastCodeML (Valle et al. 2014) starting with M1-estimated (κ and ω_0_) starting parameters, both on the Vital-IT computer cluster.

#### Ensembl Gene ID to Entrez Gene ID conversion

We use gene sets from NCBI Biosystems (Geer et al. 2010) (see next section). Since these sets are annotated with Entrez gene IDs, whereas Selectome uses Ensembl gene IDs, we created a one-to-one Ensembl–Entrez conversion table, to map the Ensembl gene IDs in the branch specific gene tables (*G_i_*) to Entrez gene IDs. First, we started with a gene list (*G_entrez_*) containing 20,016 protein coding human genes located either on the autosomal, X or Y chromosomes, downloaded from the NCBI Entrez Gene (Maglott et al. 2011) website (http://www.ncbi.nlm.nih.gov/gene, accessed on July 16, 2014). We further collected conversion tables (often containing one-to-many or many-to-many mappings) from HGNC (http://www.genenames.org/biomart/, accessed on July 16, 2014), NCBI (ftp://ftp.ncbi.nih.gov/gene/DATA, accessed on July 16, 2014), and Ensembl (version 70, http://jan2013.archive.ensembl.org/biomart/martview). We only kept the rows in these conversion tables that contained genes from *G_i_* and from *G_Entrez_*. Next, we merged the three tables to one, and only uniquely mapped genes (Ensembl ID – Entrez ID) were used further. For each Ensembl ID we kept the Ensembl-Entrez mapping with the highest count (in case of multiple candidates we chose randomly one) and we then repeated this procedure for each Entrez ID. With this final list of unique one-to-one Ensembl-Entrez ID mappings we translated the Ensembl genes in the *G_i_* tables to Entrez IDs, and unmapped genes were removed. The resulting 14,574, 15,026, 15,375 and 15,450 genes in the tables *G_1_*, *G_2_*, *G_3_* and *G_4_* respectively were used for further analyses.

#### Gene sets

We downloaded a list of 2,609 human gene sets of type ‘pathway’ from the NCBI Biosystems (Geer et al. 2010) database (http://www.ncbi.nlm.nih.gov/biosystems, accessed on July 16, 2014). The Biosystems database is a repository of gene sets collected from manually curated pathway databases, such as BioCyc (Caspi et al. 2014), KEGG (Kanehisa and Goto 2000; Kanehisa et al. 2014), The National Cancer Institute Pathway Interaction Database (Schaefer et al. 2009), Reactome (Croft et al. 2014) and Wikipathways (Kelder et al. 2012).

For each primate branch of interest (Homininae, Hominidae, Hominoidae and Catarrhini) (Figure 1), we excluded genes that could not be mapped to the corresponding gene list *G_i_* (see previous section), then removed gene sets with less than 10 genes, because the gene set enrichment test has low power to detect selection in small sets. We merged groups of nearly identical gene sets (i.e. sets that share 95% or more of their genes) into single gene sets, i.e. the union of all gene sets in these groups. In the text, the name if these union sets is followed by an asterisk (‘*’). To distinguish gene sets with identical names, their source database is added to their name. After the filtering process, we obtained *S_1_*=1,415, *S_2_*=1,424, *S_3_*=1,441 and *S_4_*=1,441 gene sets for the four branches to be tested for selection with the gene set enrichment analysis (Table 1; numbering according to Figure 1). Note that for each branch we use a different gene list (*G_i_*), leading to a different number of gene sets, as we condition on a minimum of ten genes per set for each branch.

### Data analysis

#### Test for polygenic selection

We used a gene set enrichment approach to test for polygenic signals of positive selection on the four primate branches Catarrhini, Hominoidae, Hominidae and Homininae. We first calculated for each gene set its SUMSTAT score, which is the sum of selection scores of genes in the set of interest (Tintle et al. 2009; Daub et al. 2013). As selection score we took the fourth-root of the ΔlnL*_g_* values (called ΔlnL4 hereafter) to ensure that the distribution of non-zero ΔlnLs is approximately Normal (Hawkins and Wixley 1986; Roux et al. 2014). This procedure also prevents extreme scoring genes from getting too much weight in the test, which would otherwise result in significant pathways mostly due to a few outlier genes. The SUMSTAT score of a gene set *s* is then simply calculated as:

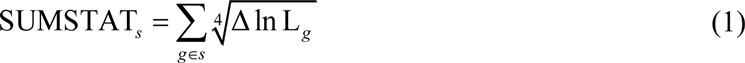

The R code to run the gene set enrichment pipeline together with detailed examples and data sets is freely available from https://github.com/CMPG/polysel.

#### Empirical null distribution

The significance of the SUSMTAT score of a pathway was inferred by creating a null distribution of random gene sets of identical size and calculating SUMSTAT on these random sets. The null distribution was built by sampling genes at random from all genes in G*_i_* that belonged to at least one gene set. To improve computation time, we created the null distribution with a sequential random sampling method (Ahrens and Dieter 1985), which avoids the burden of high precision p-value estimation for low scoring, and thus for us uninteresting, gene sets. For this we first tested all sets against a small null distribution with 10,000 random sets and estimated their p-value. For those sets with a p-value < 0.5 we expanded the null distribution with another 10,000 randomizations. This process was continued with decreasing p-value thresholds (
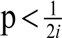 at the *i*th iteration) until we reached a maximum of 1,000,000 randomizations.

We have considered using a parametric distribution, as we know from theory and simulation studies (Zhang et al. 2005) that under the null hypothesis the log-likelihood ratios in the branchsite test should be distributed as a mixture of 50% zeros and 50% a χ^2^ distribution with one degree of freedom. Assuming such a distribution, we can infer the expected null distribution of a log-likelihood ratio score at the gene set level as well. However, in our real data the proportion of zeros is much larger, around 70%-75% for the four tested branches and we observed that a parametric null distribution produces a skewed distribution of p-values, leading to an overestimation of p-values (see also Supplemental Text S1) and thus an under-estimation of the signal for positive selection. Therefore, we have favored the use of an empirical distribution. Note that a simulation study by Yang and dos Reis (2011) produced similar high proportions of zeroes, but only for short sequence lengths (<50 codons). As only a few genes (9) in our study have such a short length, we conclude that sequence length cannot explain the high number of zero log-likelihood ratios. It would be worth investigating if other model assumptions do not fully match reality and could therefore be responsible for the deviation from the theoretical distribution of log-likelihood ratios, but this lies beyond the scope of the present study.

#### Removing sets with outlier genes

Gene sets can potentially have a high SUMSTAT score due solely to one gene with an extremely high ΔlnL value. However, we are interested here in gene sets affected by polygenic selection, where multiple genes have moderately high selection scores. Therefore, we also tested the gene sets after removing their highest scoring gene, and contrasted their SUMSTAT score against random sets that also had their top scoring gene removed. Those gene sets that were not scoring significant anymore after this test were not included in further analyses.

#### Removing redundancy in overlapping gene sets (‘pruning’)

The gene set enrichment test can result in partially redundant gene sets being called significant, because they share high scoring genes, and BioSystems includes overlapping or redundant sets. We therefore removed the overlap between gene sets with a ‘pruning’ method similar to one we described in a previous study (Daub et al. 2013). In short, we removed for each branch the genes of the most significant pathway from all the other pathways, and ran the enrichment test on these updated gene sets. We repeated this pruning procedure until no sets were left to be tested.

We estimated the False Discovery Rate (FDR) in our results empirically, since the tests in the pruning procedure are not independent and the results are biased toward low p-values (only the high scoring sets will remain after pruning). To estimate the FDR, we repeatedly (*N*=300) permuted ΔlnL4 scores among genes that are part of a set, and tested the gene sets by applying the above described pruning method. For each observed p-value *p\** in our original results, we can estimate the FDR (if we would reject all hypotheses with a p-value ≤ *p\**) with:

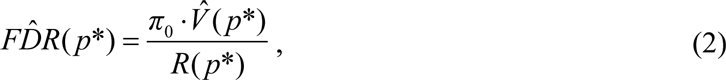

where *π*_0_ is the proportion of true null hypotheses, 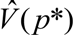 is the estimated number of rejected true null hypotheses if all hypotheses are true nulls and *R*(*p\**) is the total number of rejected hypotheses. We conservatively set *π*_0_ = 1, and estimated 
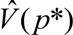 from the mean proportion of gene sets in the randomized data sets with p-value ≤ *p*^*^. The q-value was finally determined by taking the lowest estimated FDR among all observed p-values ≥ *p*\*. We reported the gene sets that scored significant (q-value < 0.2) both before and after pruning.

#### Classical GO enrichment test

To contrast our results with a classical Gene Ontology (GO) enrichment test, we defined for each branch a list of top scoring genes, namely those genes with a ΔlnL value resulting in a q-value < 0.2 when compared against a null distribution consisting of 50% zeros and 50% χ^2^ with one degree of freedom (see paragraph ‘Empirical null distribution’ and Zhang et al. 2005). Note that this is a much less conservative score compared to the branches reported on the Selectome website, as the latter are based on q < 0.10 and were computed over all branches and trees using a more conservative null distribution (χ^2^ with one degree of freedom). The top genes thus defined were used as input for the online tool Fatigo (http://babelomics.bioinfo.cipf.es, (Al-Shahrour et al. 2004)). We tested for enrichment in (level 3-9) GO biological processes and molecular functions using as background the gene lists *G_i_*. The resulting p-values were corrected for multiple testing by calculating the q-value per branch and GO category (biological process and molecular function), where q-values < 0.2 were considered significant.

#### Test for bias in genes filtered from Selectome

Since all gene trees with less than six leaves after removing unreliably aligned sequences were excluded in Selectome, we tested which categories of gene sets where enriched with these excluded genes, and were thus unlikely to show significant results in our gene set enrichment test. For this we performed for each gene set a Fisher’s exact test on a contingency table with the counts of included and excluded genes in the set contrasted against the same counts for the rest of the genes in *G_entrez_*, the list of genes downloaded from the NCBI Entrez website. The resulting p-values were corrected for multiple tests, and gene sets with a q-value < 0.2 were reported.

#### Investigating bias in branch length and GC content

To study whether branch length or GC content might have biased our results, we calculated the correlation between the selection score ΔlnL4 and each of these factors. The (gene specific) synonymous branch length, dS, was obtained by running the *M1* model in *codeml* with branch length optimization. Due to time constraints, the largest gene tree (ENSGT00550000074383, subtree 1) was excluded from the calculations. GC content was computed at two different levels. At the branch level, GC content was estimated using the tool *nhphyml* (Boussau and Gouy 2006), with settings: branch length optimization, no tree topology optimization, an infinite number of GC categories, 4 gamma rate categories (alpha estimated) and nucleotide mode. At the gene tree level, the average, minimum and maximum GC content together with the variance in GC content were calculated. At both levels the unmasked sequences from the terminal branches in each gene tree served as input for the computations.

#### Correcting for multiple testing

For all tests (except when inferring significance after pruning), we calculated the q-value (Storey J. D. and Tibshirani 2003; Storey John D. et al. 2004) as a measure of the false discovery rate using the R package *qvalue* (with parameter *pi0.method* set to “bootstrap”) and reported those gene sets with a q-value < 0.2.

#### Enrichment maps

The enrichment maps (Supplemental Figures S2 to S5 and S7) were created in Cytoscape v. 2.8.3 with the *Enrichmentmap* plugin (Merico et al. 2010).

## Acknowledgements

This work was supported by the Swiss National Science Foundation (grant numbers PDFMP3-130309 to LE and 31003A_153341 and CR32I3_143768 to MRR). The computations were performed at the Vital-IT (http://www.vital-it.ch) Center for high-performance computing of the SIB Swiss Institute of Bioinformatics, on the Ubelix HPC cluster of the University of Bern and on the Swiss Multi-Science Computing Grid.

## Text S1

### Using a parametric null distribution

If we assume independence between genes in a set, we can define the gene set level likelihoods, LH0*_s_* and LH1*_s_*, as the product of the gene level likelihoods:

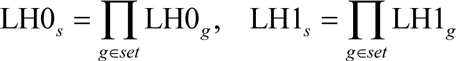

We can then calculate a gene set level likelihood ratio test score (ΔlnL*_s_*), which is equal to the sum of ΔlnL*_g_* scores of genes in the gene set:

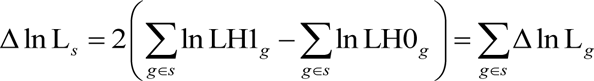

This ΔlnL*_s_* score can then be compared against a parametric null distribution to infer its significance.

In the branch-site test for positive selection, the dN/dS ratio for the class of sites under positive selection (ω_2_) is constrained by ω_2_ ≥ 1 and the null model H_0_ is nested in H_1_ with ω_2_ fixed on the boundary of the parameter space (ω_2_ = 1). Therefore, in case of no positive selection (H_0_ is true), we would expect that the ΔlnL values at the gene level (ΔlnL*_g_*) are distributed as a mixture of 50% zeros and 50% (Self and Liang 1987; Zhang et al. 2005). For the gene set level ΔlnL values (ΔlnL*_s_*), we have *N* free parameters for a gene set of size *N*. The expected null distribution of ΔlnL*_s_* would be a mixture of χ^2^ distributions with 0, 1, …, *N* degrees of freedom, where the mixture weights are the binomial coefficients (Ota et al. 2000). For example, with gene sets containing three genes, the distribution would be:

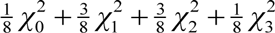

The probability for observing a ΔlnL*_s_* of *x* or higher in gene sets of size three can then be calculated as:

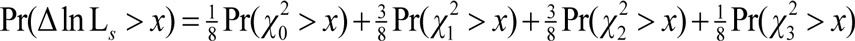

The ΔlnL_*s*_ scores that we use have a higher proportion of zero’s, namely around 70-80%. Testing our gene sets against a parametric null distribution, results therefore in unacceptable p-value distributions, which is shown by the following qq-plots comparing the p-values from a parametric test with an expected uniform distribution:

**Table S1.**
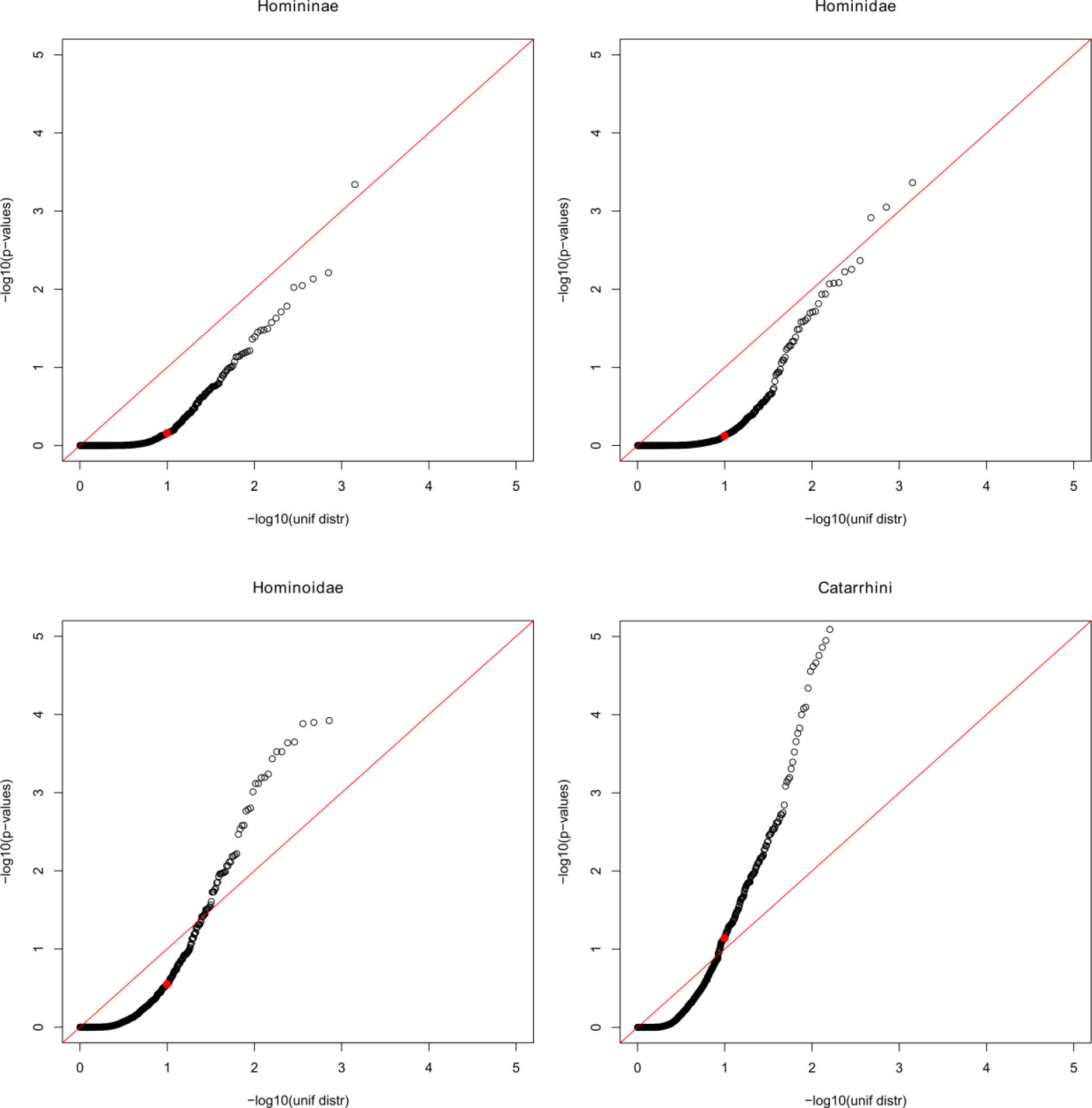
This table is a separate excel file (TableS1.xlsx) which contains three tables: (A) Top scoring gene sets in the gene set enrichment test before pruning, their rank and score and p- and q-value (also after leaving out the highest scoring gene). All gene sets with q<0.2 are reported. (B) All significantly scoring pathways and their rank (before pruning) in all four branches. An ‘-’ indicates that the set was not defined in that particular branch. Gene sets that scored significant after pruning are marked in bold. (C) Genes in top scoring gene sets and their ΔlnL4 and ΔlnL scores.

**Table S2.** 

**A.** Correlation (Pearson’s r) between selection score ΔlnL4 and GC statistics at the gene level

| Per branch | $r$ | [ci1, ci2] | p-value |
| --- | --- | --- | --- |
| GC content | -0.045 | [-0.054, -0.036] | <2.2e-16 |
| Per tree | $r$ | [ci1, ci2] | p-value |
| mean GC content | -0.043 | [-0.052, -0.034] | <2.2e-16 |
| GC variance | 0.044 | [0.036, 0.053] | <2.2e-16 |
| max GC – min GC | 0.074 | [0.065, 0.083] | <2.2e-16 |

[ci1, ci2]: 95% confidence interval

**B.** Correlation (Pearson’s r) between gene set enrichment score (-log[p-value]) and GC statistics at the gene set level. Significant correlations with |r| > 0.1 are marked in bold.

| Per branch |  |  |  |
| --- | --- | --- | --- |
| GC content | $r$ | [ci1, ci2] | p-value |
| Homininae | -0.050 | [-0.102, 0.002] | 0.058 |
| Hominidae | -0.068 | [-0.121, -0.016] | 0.011 |
| Hominoidae | 0.013 | [-0.041, 0.066] | 0.638 |
| Catarrhini | -0.040 | [-0.094, 0.014] | 0.144 |
| Per tree |  |  |  |
| mean GC content | $r$ | [ci1, ci2] | p-value |
| Homininae | -0.034 | [-0.086, 0.018] | 0.202 |
| Hominidae | -0.066 | [-0.118, -0.014] | 0.013 |
| Hominoidae | 0.022 | [-0.03, 0.074] | 0.402 |
| Catarrhini | -0.021 | [-0.073, 0.03] | 0.417 |
| GC variance | $r$ | [ci1, ci2] | p-value |
| Homininae | **0.183** | [0.132, 0.233] | 3.83e-12 |
| Hominidae | **0.172** | [0.121, 0.222] | 5.87e-11 |
| Hominoidae | **0.154** | [0.103, 0.204] | 4.46e-09 |
| Catarrhini | **0.186** | [0.135, 0.235] | 1.18e-12 |
| max GC – min GC | $r$ | [ci1, ci2] | p-value |
| Homininae | 0.033 | [-0.019, 0.085] | 0.217 |
| Hominidae | -0.007 | [-0.059, 0.045] | 0.803 |
| Hominoidae | -0.063 | [-0.114, -0.011] | 0.017 |
| Catarrhini | 0.018 | [-0.034, 0.07] | 0.493 |

[ci1, ci2]: 95% confidence interval

Table S3 This table is a separate excel file (TableS3.xlsx) comparing results of the gene set enrichment test on a dataset of gene trees with or without duplications. It contains five tables: The number of available genes and gene sets for each branch excluding or including duplications (A), and -for each of the four tested branches- top scoring gene sets in the enrichment test with duplications and their score in the test without duplications (B-E).

Table S4 This table is a separate excel file (TableS4.xlsx) presenting results of the gene set enrichment test where missing branches were not given the ΔlnL4 score of their child branch. It contains three tables: The number of available genes and gene sets for each branch excluding or including missing branches (using the ΔlnL4 score of their child branch) (A), and two tables with top scoring gene sets in the enrichment test before (B) and after (C) pruning, their rank and score and p- and q-value. All gene sets with q-value<0.2 are reported.

**Figure S1.**
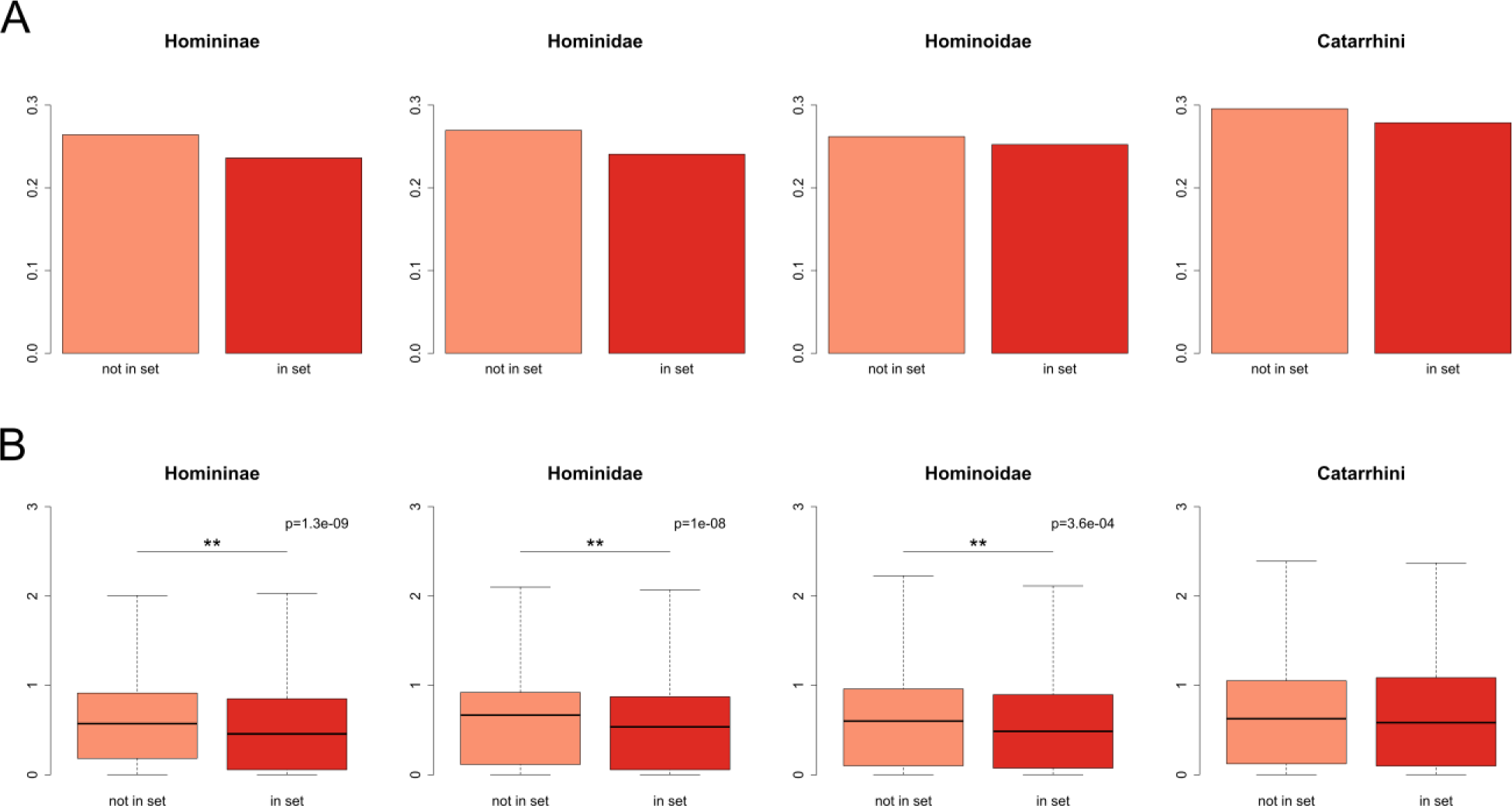
Proportion (A) and distribution (B) of non-zero ΔlnL4 scores in the genes in the four tested branches, split between genes that are not part of a gene set and genes that are part of at least one gene set. Especially the Homininae and Hominidae branches show a lower proportion and lower values of non-zero scores for genes in sets. The significance of the difference in distributions of non-zero ΔlnL4 scores is tested with a Wilcoxon signed-rank test.

**Figure S2.**
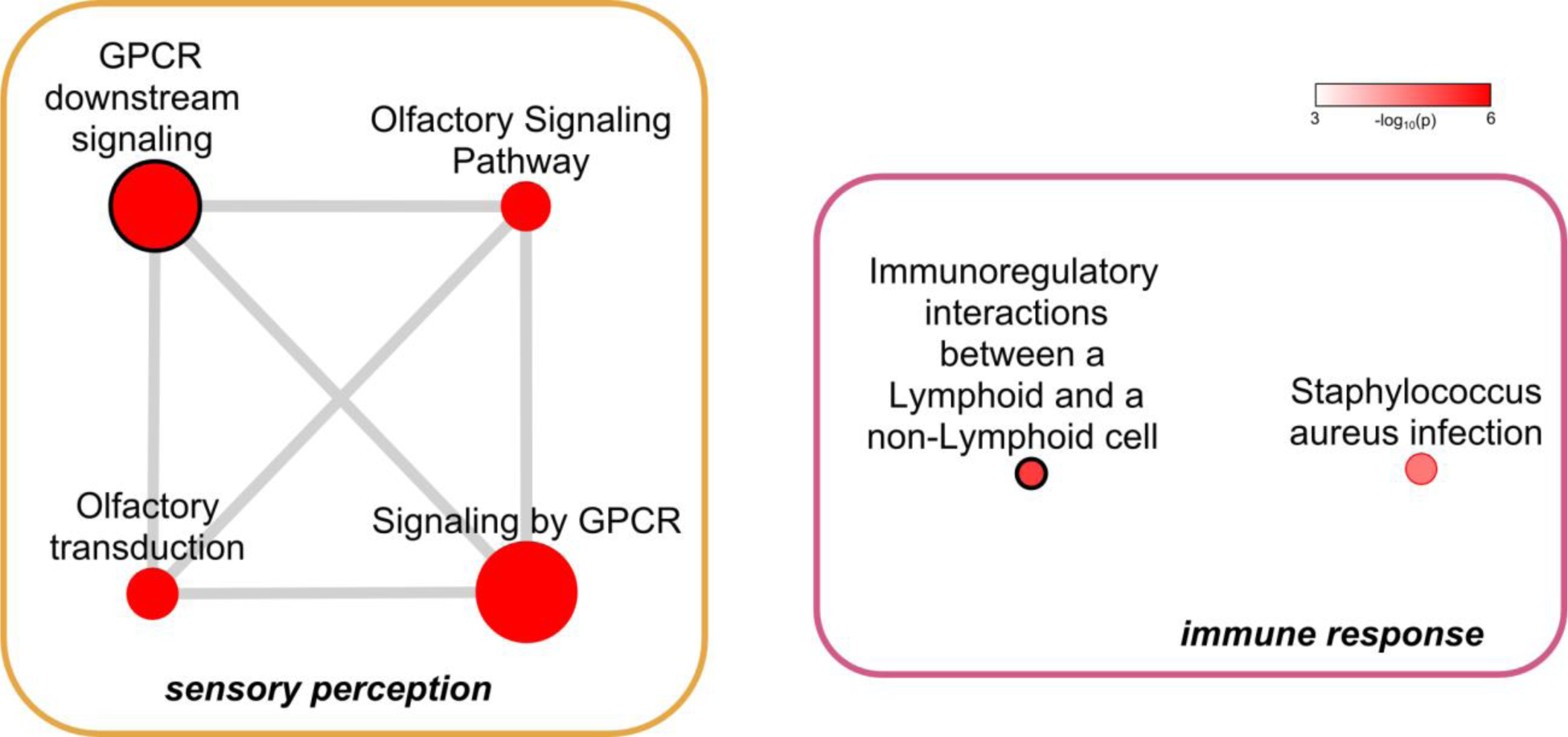
Enrichment map of the gene sets enriched for signals of positive selection in the Homininae branch. The nodes represent the 6 pathways with q-values < 0.2 before pruning (removal of overlapping genes). The node color scale represents gene set p-values. Edges represent mutual overlap; nodes are connected if one of the sets has at least 33% of its genes in common with the other gene set. The widths of the edges scale with the similarity between nodes. The two pathways that remained significant after pruning are marked with a black circle. A full-size version of this figure is available at https://dx.doi.org/10.6084/m9.figshare.3119029.

**Figure S3.**
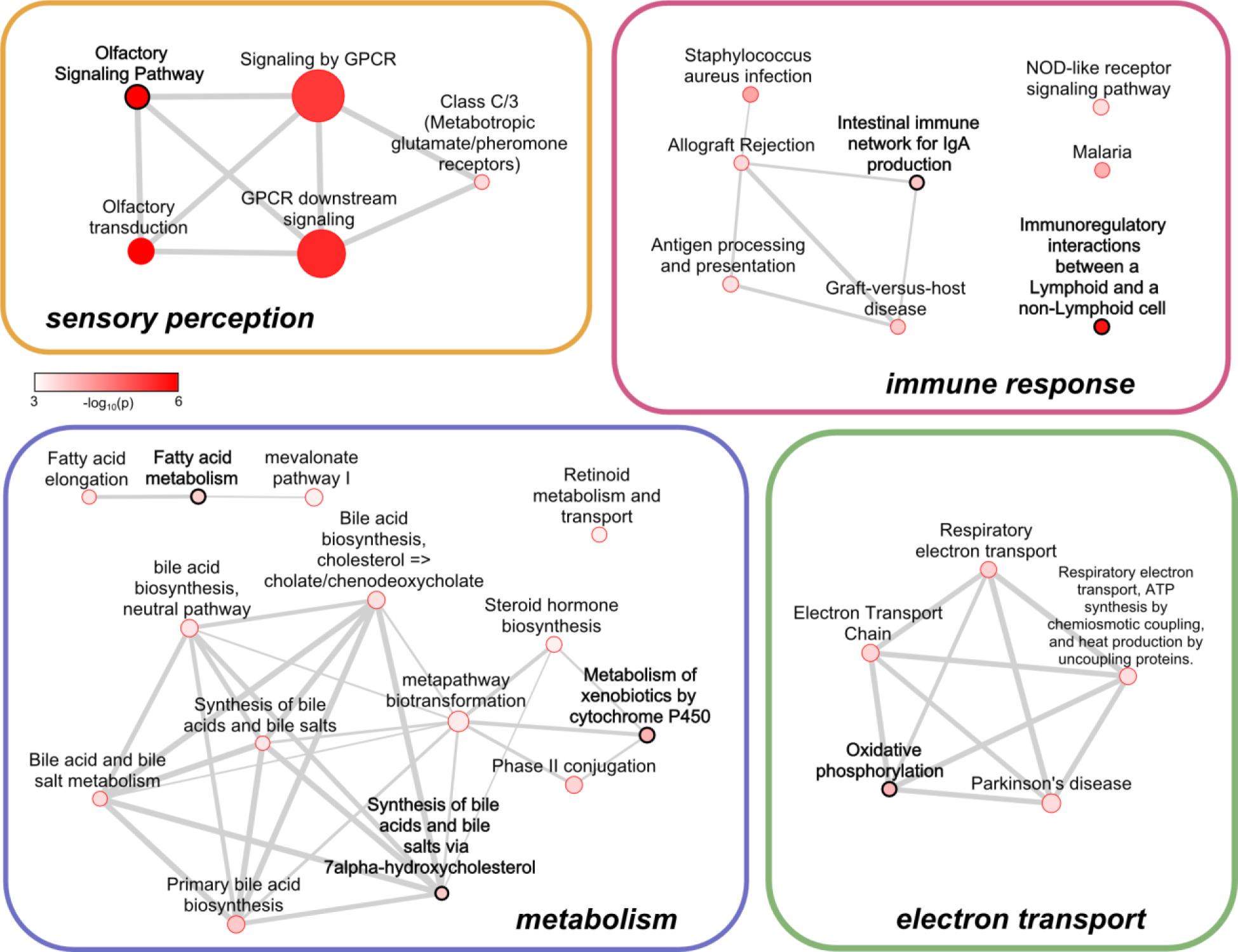
Enrichment map of the gene sets enriched for signals of positive selection in the Hominidae branch. The nodes represent the 32 pathways with q-values < 0.2 before pruning (removal of overlapping genes). The seven pathways that remained significant after pruning are marked with a black circle. See Figure S2 for a more detailed explanation of the enrichment map. Nodes marked with * represent unions of pathways that share more than 95% of their genes. A full-size version of this figure is available at https://dx.doi.org/10.6084/m9.figshare.3119029.

**Figure S4.**
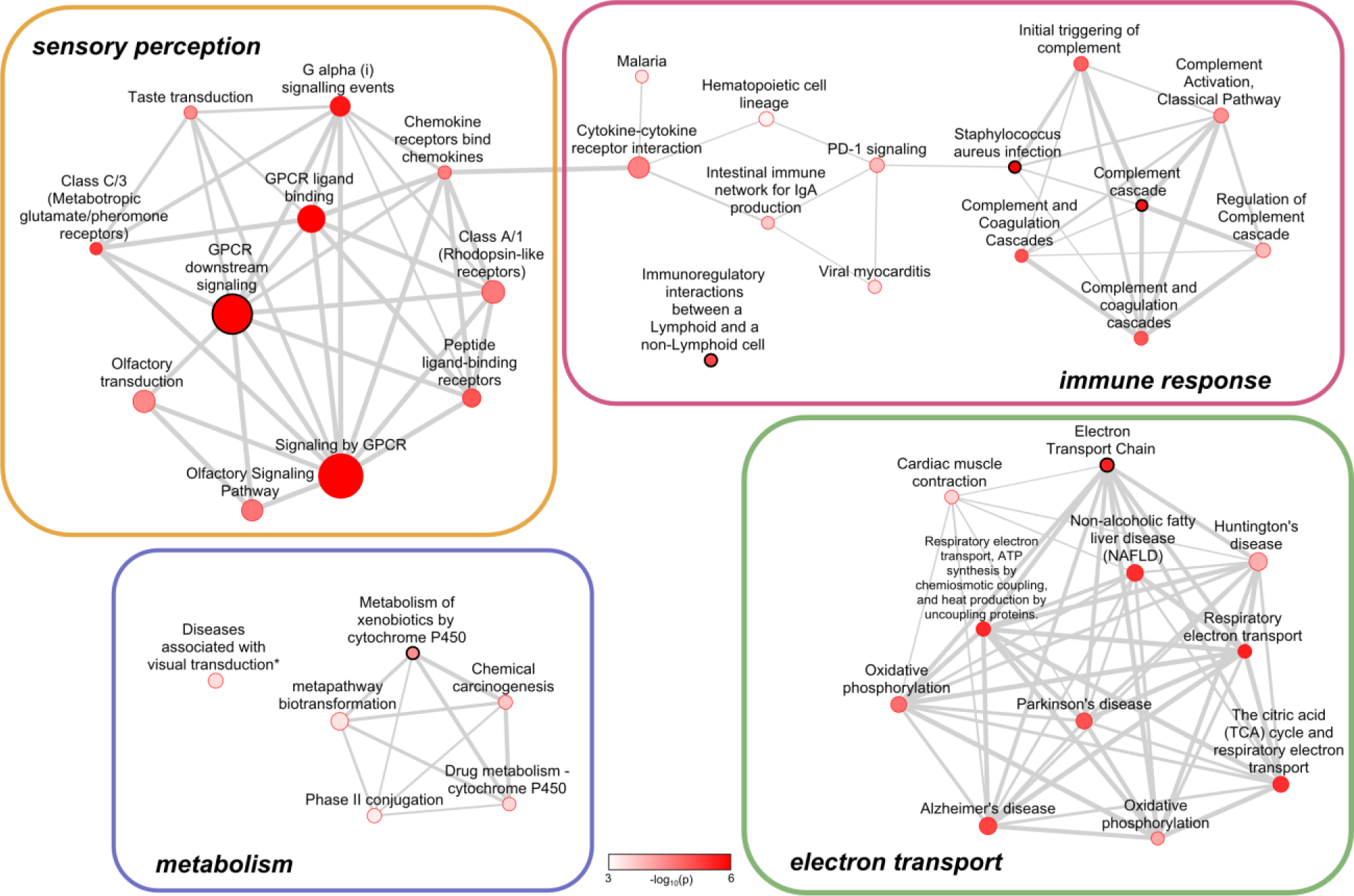
Enrichment map of the gene sets enriched for signals of positive selection in the Hominoidae branch. The nodes represent the 42 pathways with q-values < 0.2 before pruning (removal of overlapping genes). The six pathways that remained significant after pruning are marked with a black circle. See Figure S2 for a more detailed explanation of the enrichment map. A full-size version of this figure is available at https://dx.doi.org/10.6084/m9.figshare.3119029.

**Figure S5.**
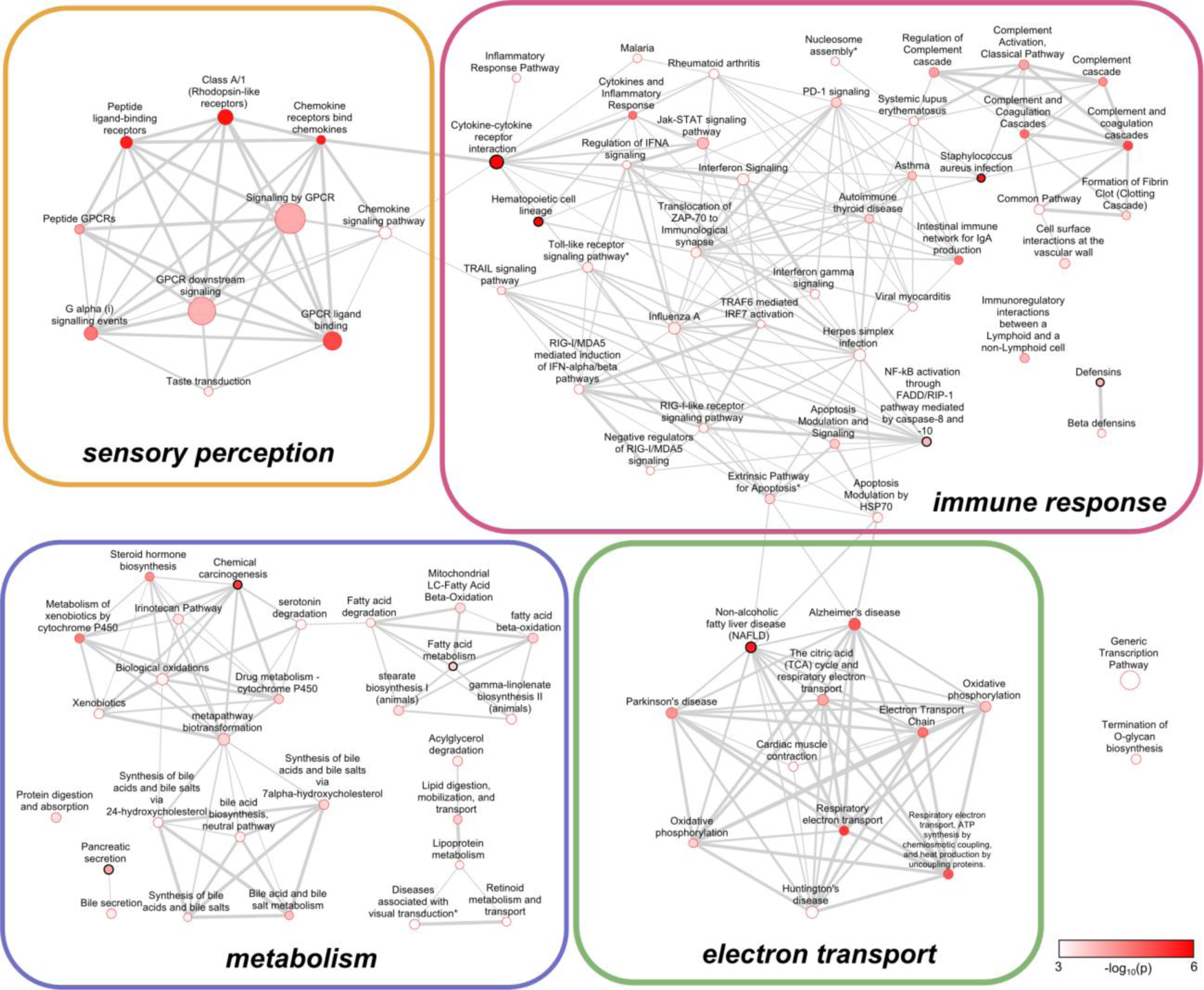
Enrichment map of the gene sets enriched for signals of positive selection in the Catarrhini branch. The nodes represent the 93 pathways with q-values < 0.2 before pruning (removal of overlapping genes). The nine pathways that remained significant after pruning are marked with a black circle. See Figure S2 for a more detailed explanation of the enrichment map. Nodes marked with * represent unions of pathways that share more than 95% of their genes. A full-size version of this figure is available at https://dx.doi.org/10.6084/m9.figshare.3119029.

**Figure S6.**
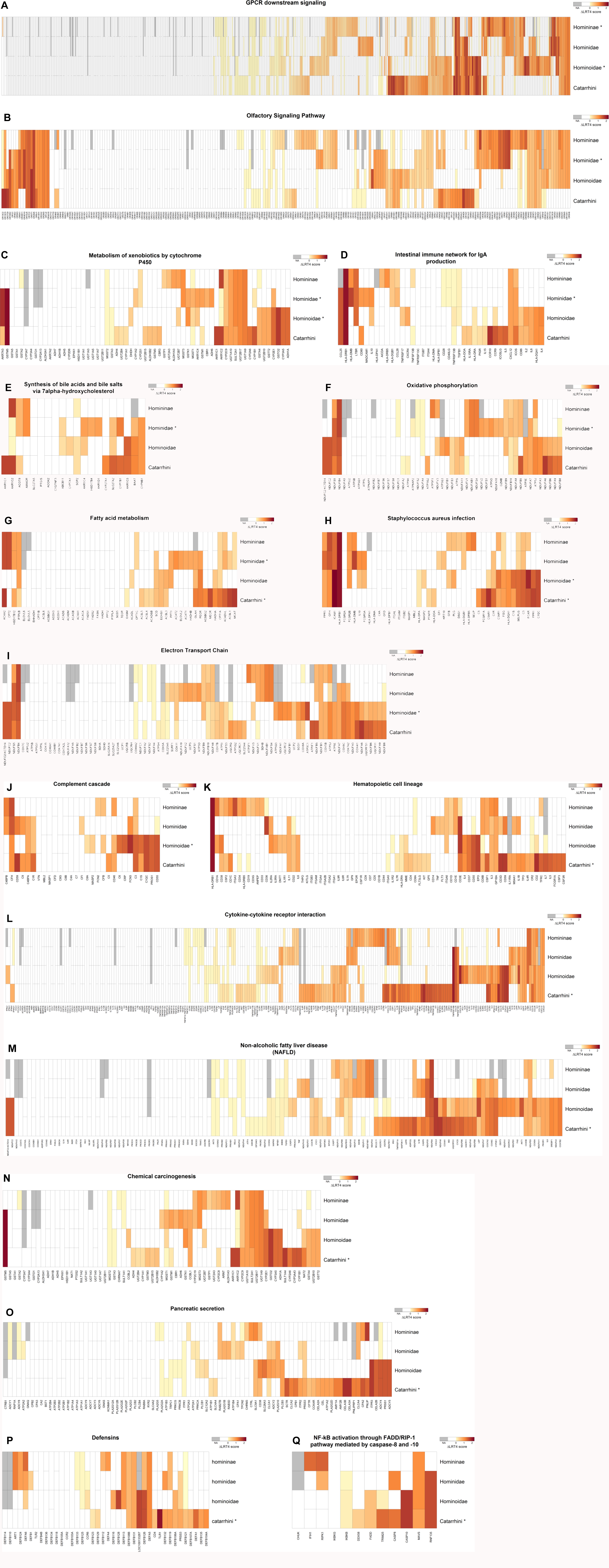
Heat maps showing branch specific ΔlnL4 scores of genes in pathways that score significant (q < 0.2) in the gene set enrichment test after pruning. The ΔlnL4 score is computed as the fourth root of the log-likelihood ratio in the branch site test for positive selection. Branches where a pathway scores significant are marked with a ‘*’. The genes are grouped by hierarchical clustering to visualize blocks with similar signals among branches. Genes for which ΔlnL4 scores were not available (NA) in a certain branch are depicted in grey. Genes are merged (horizontally) with their paralog(s) into an ‘ancestral gene’ in the branches preceding a duplication and their scores were included only once in the calculation of the SUMSTAT score for these branches. Genes with (vertically) merged branches represent cases where the sequence of one or more species is missing or excluded, resulting in a single ‘average’ ΔlnL4 score over multiple branches. We used this score when testing each branch separately. Full-size versions of these heat maps are available at https://dx.doi.org/10.6084/m9.figshare.3119026.

**Figure S7.**
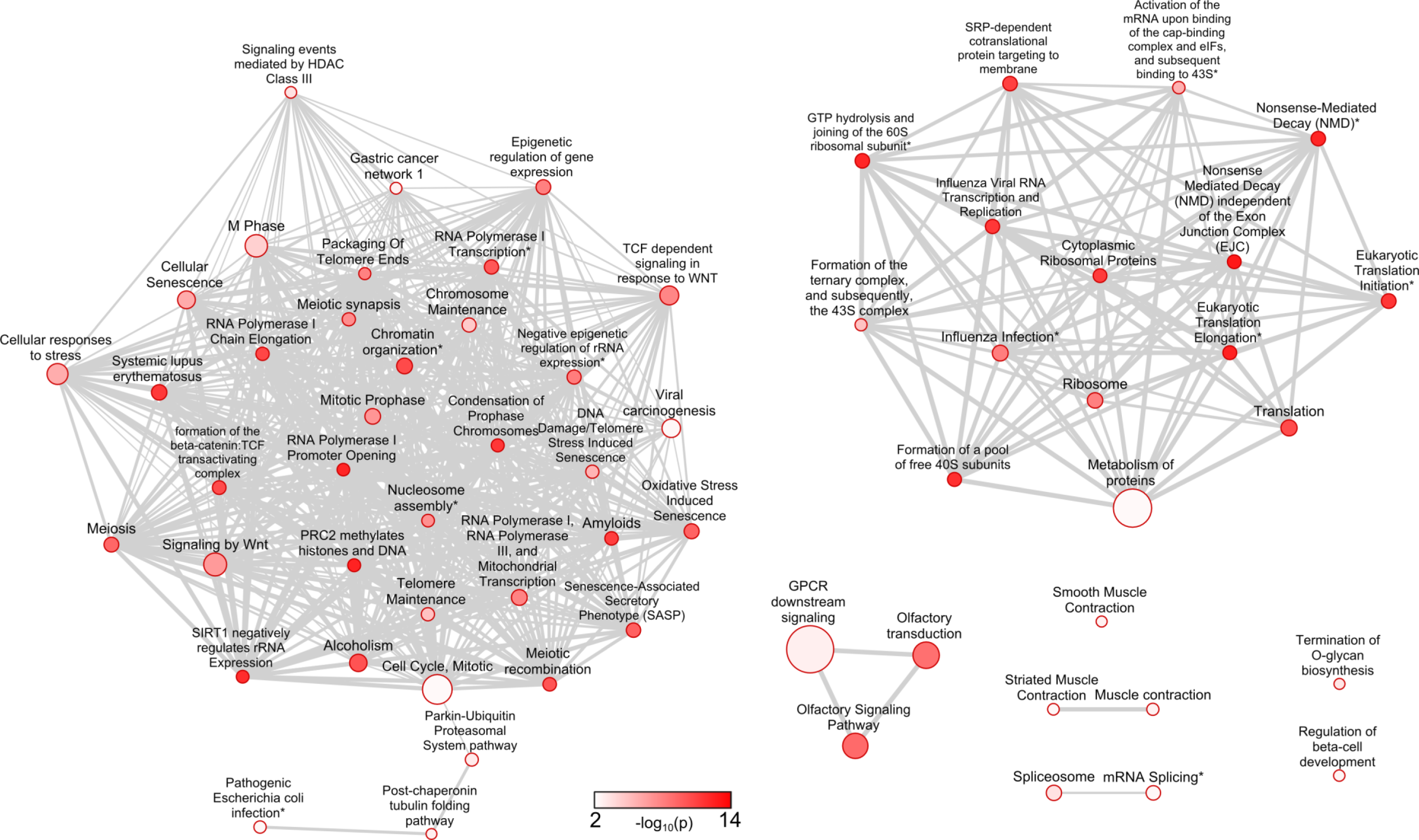
Enrichment map of gene sets enriched with genes that were excluded in Selectome. The nodes represent the 62 pathways with q-values < 0.2 in a Fisher’s exact test. See Figure S2 for a more detailed description of the enrichment map. Nodes marked with * represent unions of pathways that share more than 95% of their genes. A full-size version of this figure is available at https://dx.doi.org/10.6084/m9.figshare.3119122.

**Figure S8.**
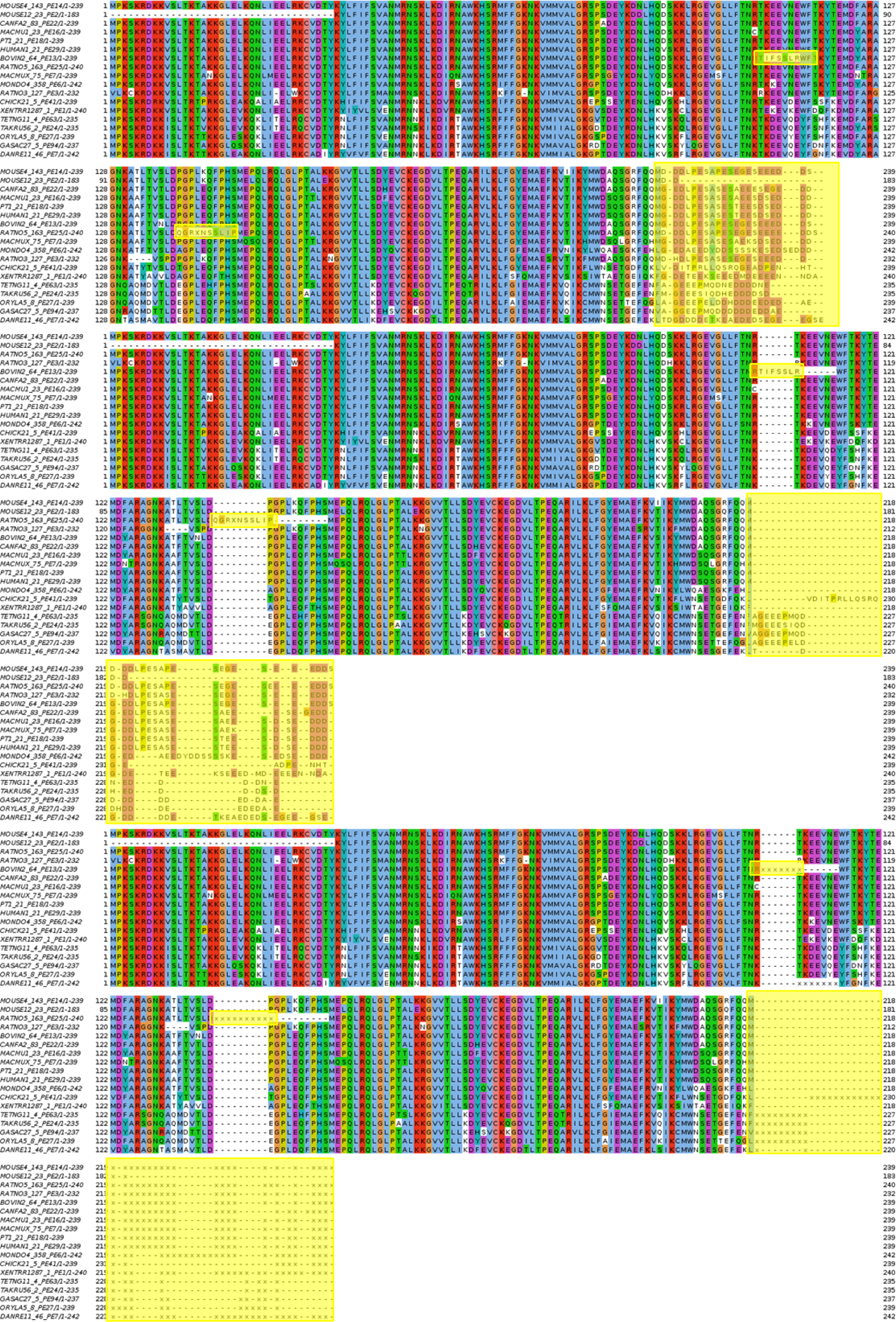
MSA filtering example. a) Before alignment with PAGAN. Problematic regions are highlighted in yellow (2 non-homologous regions and one region with repeats). b) After alignment with PAGAN. Non-homologous regions are almost entirely isolated from the rest. c) End of the pipeline. Non-homologous or difficult to align regions are masked with ‘x’, including the region containing repeats.

**Figure S9.**
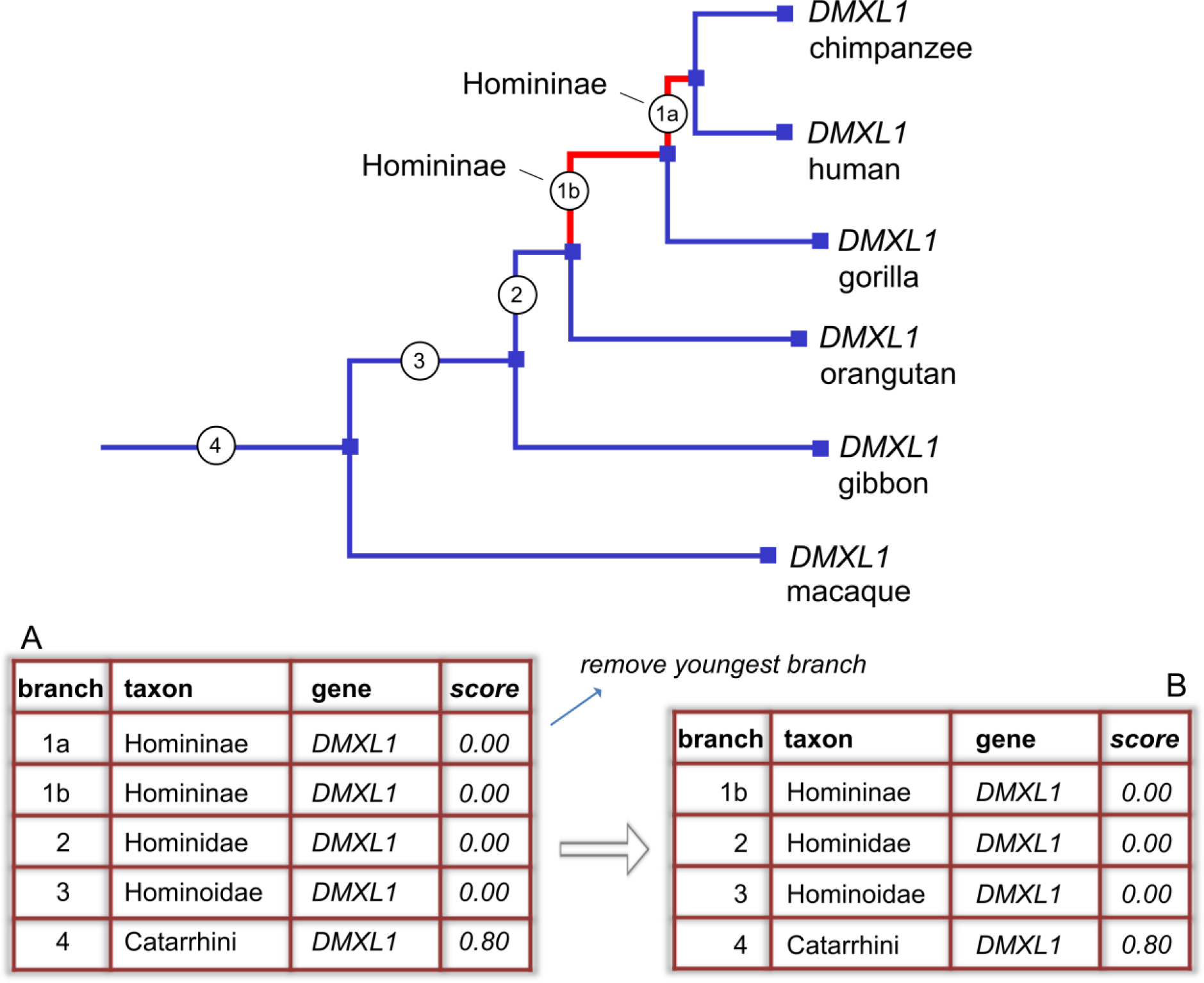
Example of multiple Homininae branches (branch *1a* and *1b*, marked red) in the *DMXL1* gene tree. Only the oldest branch (*1b*) will be used in the enrichment test. Table A illustrates that initially the Homininae branches occur in double in the table, while after removing the youngest Homininae branch, only one row per branch (taxon) and human gene remains (Table B). It should be emphasized that this is not a biological phenomenon, but simply a technical result of the labeling choices made in the Ensembl database, which are carried into the Selectome database and thus into the dataset used in this study.

**Figure S10.**
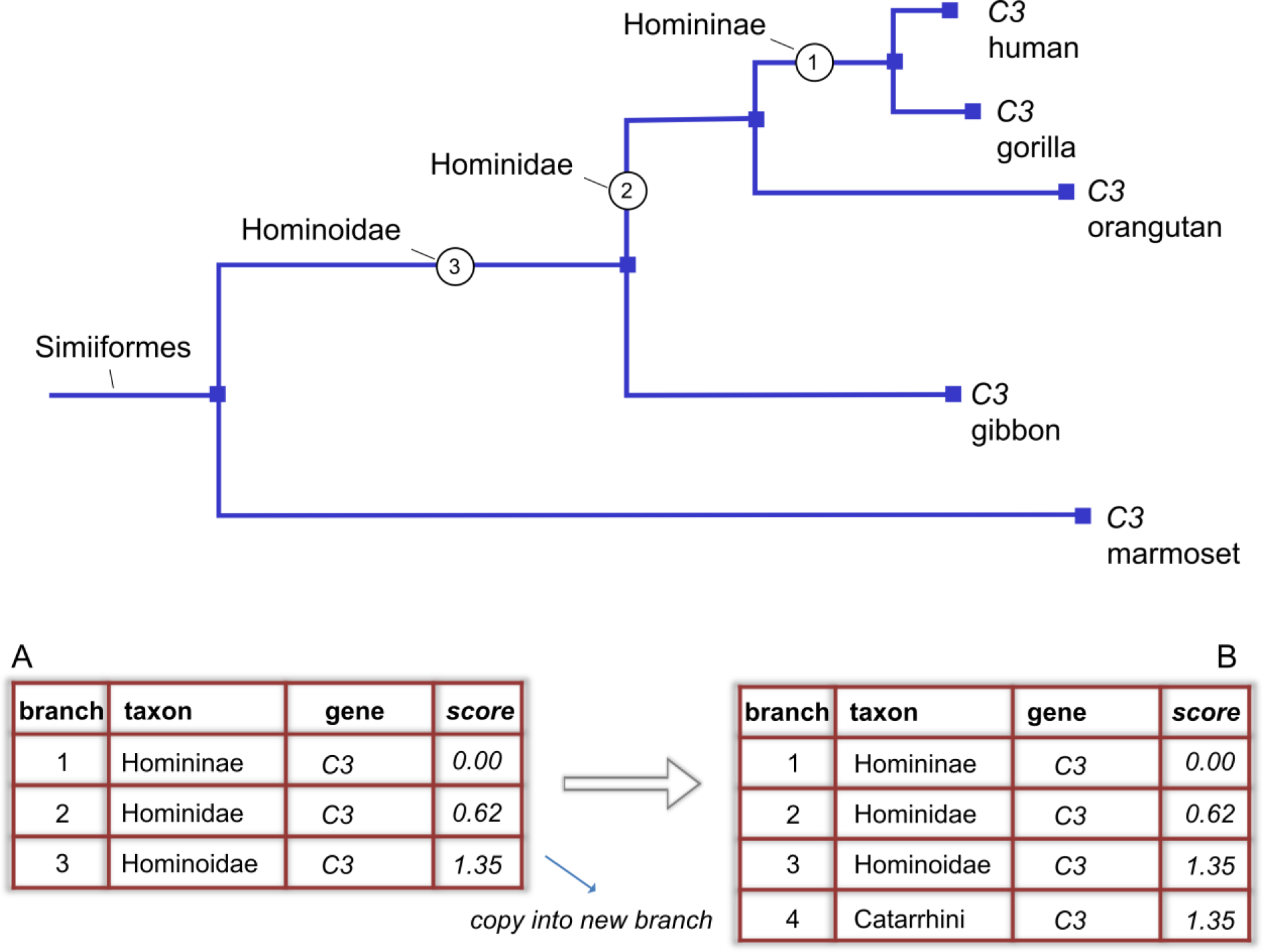
Example of a missing branch (Catarrhini) in the CE3 gene tree, due to an absent or low quality macaque sequence for this gene. The score assigned to the Hominoidae branch actually represents an ‘average’ score for both the Hominoidae and Catarrhini branch, and therefore we use this score for both branches in the enrichment test. Table A illustrates that initially one branch is missing, while after copying the Hominoidae score into a new Catarrhini row, one row per branch (taxon) and human gene is now available (Table B).

**Figure S11.**
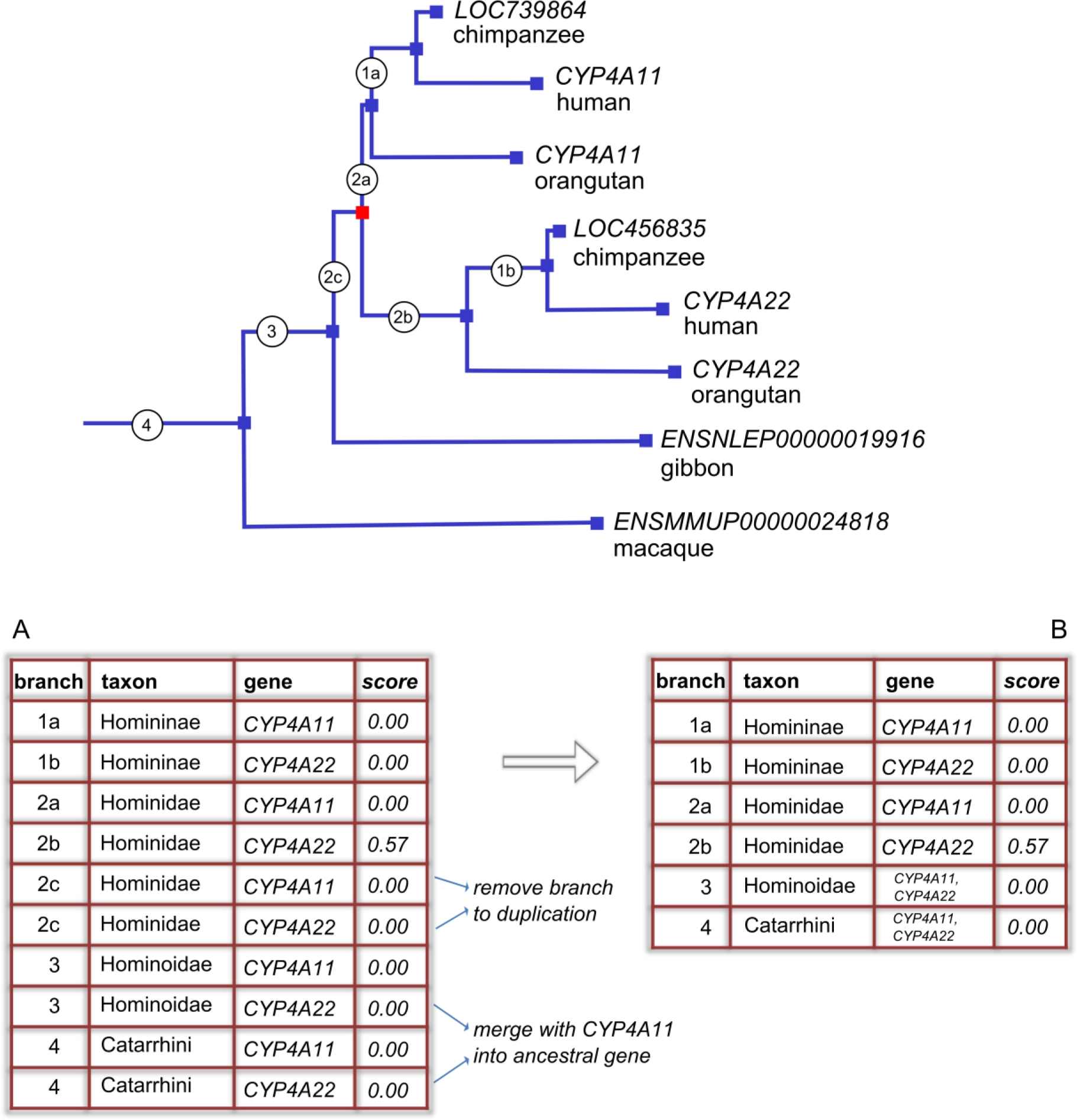
Example of a duplication event (red node) in the Hominidae branch of the *CYP4A11*/*CYP4A22* gene tree. In the enrichment tests (i) the branch leading to the duplication (*2c*) is removed and (ii) in the Hominoidae (*3*) and Catarrhini branch (*4*), the ‘ancestral’ gene *CYP4A11, CYP4A22* replaces the two human paralogs *CYP4A11* and *CYP4A22*. Table A illustrates that initially some branches (and their scores) occur in double in the table. After merging and removing branches, only one row per branch (taxon) and human gene remains (Table B).

